# A hierarchical watershed model of fluid intelligence in childhood and adolescence

**DOI:** 10.1101/435719

**Authors:** D. Fuhrmann, I. L. Simpson-Kent, J. Bathelt, the CALM team, R. A. Kievit

## Abstract

Fluid intelligence is the capacity to solve novel problems in the absence of task-specific knowledge, and is highly predictive of outcomes like educational attainment and psychopathology. Here, we modelled the neurocognitive architecture of fluid intelligence in two cohorts: CALM (*N* = 551, aged 5 - 17 years) and NKI-RS (*N* = 335, aged 6 - 17 years). We used multivariate Structural Equation Modelling to test a preregistered watershed model of fluid intelligence. This model predicts that white matter contributes to intermediate cognitive phenotypes, like working memory and processing speed, which, in turn, contribute to fluid intelligence. We found that this model performed well for both samples and explained large amounts of variance in fluid intelligence (*R*^2^_CALM_ = 51.2%, *R*^2^_NKI-RS_ = 78.3%). The relationship between cognitive abilities and white matter differed with age, showing a dip in strength around ages 7 - 12 years. This age-effect may reflect a reorganization of the neurocognitive architecture around pre- and early puberty. Overall, these findings highlight that intelligence is part of a complex hierarchical system of partially independent effects.

---

Fluid intelligence (*g*_f_) is a core part of human cognition and refers to the capacity to solve novel problems in the absence of task-specific knowledge. It is highly predictive of a number of important life span outcomes, including educational attainment (Primi et al. 2010; Roth et al. 2015) and psychopathology (Gale et al. 2010). Despite years of investigation, however, our understanding of the neurocognitive architecture of *g*_f_ remains limited. Longstanding debates have considered, for instance, how *g*_f_ relates to more fundamental cognitive functions such as working memory and processing speed, and how all of these cognitive functions relate to brain structure and function (Kyllonen and Christal 1990; Fry and Hale 2000; Chuderski 2013; Ferrer et al. 2013).

Working memory is the ability to hold and manipulate information in the mind short-term. It has been suggested that working memory is a key determinant of *g*_f_ by limiting mental information processing capacity (Fukuda et al. 2010; Chuderski 2013). Proponents of this working memory account of *g*_f_ cite high correlations between the two domains ranging from 0.5 to 0.9 in meta-analyses (Ackerman et al. 2005; Oberauer et al. 2005). Such high correlations have led some to suggest that *g*_f_ and working memory are, in fact, isomorphic (Kyllonen and Christal 1990). However, more recent work has highlighted that this isomorphism only arises under conditions of high time constraints for *g*_f_ tasks (Chuderski 2013). This suggests that *g*_f_ and working memory are, in fact, separable constructs and underlines the importance of processing speed for *g*_f_.

Processing speed, the speed of mental computations, is thought to be rate-limiting to *g*_f_ and is therefore sometimes proposed to be a particularly good predictor of *g*_f_ (Kail and Salthouse 1994; Salthouse 1996; Ferrer et al. 2013; Kail et al. 2015; Schubert et al. 2017). Proponents of the processing speed account of *g*_f_ cite moderate but robust correlations between *g*_f_ and processing speed of 0.2 in meta-analyses (Sheppard and Vernon 2008) as well as longitudinal evidence (Finkel et al. 2005; Coyle et al. 2011; Kail et al. 2015). Salthouse (1996) argued in the context of cognitive aging, that processing speed determines high-level cognitive performance because slow processing means that relevant sub-operations cannot be completed in a set amount of time or are not available for successful integration. A complementary explanation of individual differences in *g*_f_ proposes that processing speed may be a direct reflection of fundamental neuroarchitectonic properties of the brain, such as myelination or white matter microstructure (Lu et al. 2011; Chevalier et al. 2015).

White matter shows protracted development throughout childhood and adolescence, and into the third decade of life (Mills et al. 2016). White matter tracts can be characterised *in vivo* using diffusion-tensor imaging (DTI), which is sensitive, but not necessarily specific, to white matter microstructural properties such as myelination or axonal density (Jones et al. 2013; Wandell 2016). Fractional anisotropy (FA) is the most commonly investigated DTI measure and quantifies the directionality of water diffusion in different white matter tracts (Pfefferbaum et al. 2000; Wandell 2016). Working memory, processing speed and *g*_f_ have each been linked to individual differences in FA (Vestergaard et al. 2011; Kievit, Davis, Griffiths, Correia, CamCAN, et al. 2016; Bathelt et al. 2018). While some studies, using Principal Component Analysis, have posited that FA in different tracts can be summarized by sizable single components (Penke et al. 2010; Cox et al. 2016), formal investigations using confirmatory factor analysis have demonstrated that single-factor models of FA generally show poor fit and do not adequately capture individual differences in white matter microstructure (Lövdén et al. 2013; Kievit, Davis, Griffiths, Correia, Cam-CAN, et al. 2016). In a similar vein, there is a growing body of literature showing specific associations between white matter tracts and cognitive abilities, with those connecting frontoparietal regions usually showing largest contributions to complex cognitive functions like *g*_f_ (Vestergaard et al. 2011; Kievit et al. 2016; Bathelt et al. 2018).

We here seek to address several critical outstanding issues in the field: First, there is limited systematic evidence on the concurrent relationships between *g*_f_, working memory, processing speed and white matter. This leaves the relative contributions of processing speed and working memory to *g*_f_ unclear, which, in turn, poses challenges for the design of effective cognitive training interventions. Second, studies usually use a single task as a proxy for complex and abstract constructs such as processing speed, working memory, and *g*_f_. This raises questions about the generalizability of findings (Noack et al. 2014). Third, our understanding of how the relationships between relevant cognitive domains and between brain and cognition change with age remains limited, raising the possibility that brain-behaviour relationships may change with age (Garrett 1946; Johnson 2000; Tamnes et al. 2017).

To address these issues, we here used structural equation modelling (SEM) to model the associations between *g*_f_, working memory, processing speed, and white matter microstructure and age in two large, independent samples: the Centre for Attention, Leaning and Memory sample (CALM, *N* = 551, aged 5 - 17 years), which consists of children and adolescents referred to a clinic for having problems with attention, learning and memory (Holmes et al. 2018), and the Enhanced Nathan Kline Institute – Rockland Sample (NKI-RS, *N* = 335, aged 6 - 17 years), a community-ascertained sample (Nooner et al. 2012).

To investigate the neurocognitive architecture of *g*_f_ in a principled way, we used a watershed model of individual differences. Based on the metaphor of a watershed, the model predicts a hierarchical many-to-one mapping of partially independent effects such that upstream tributaries (e.g. brain structure) contribute to intermediate cognitive phenomena (cognitive endophenotypes, e.g. working memory and processing speed), which then contribute to downstream, complex cognitive phenomena such as *g*_f_ (Cannon and Keller 2006; Kievit, Davis, Griffiths, Correia, CamCAN, et al. 2016). See Figure 1 for a representation of the model.

**Figure 1.**
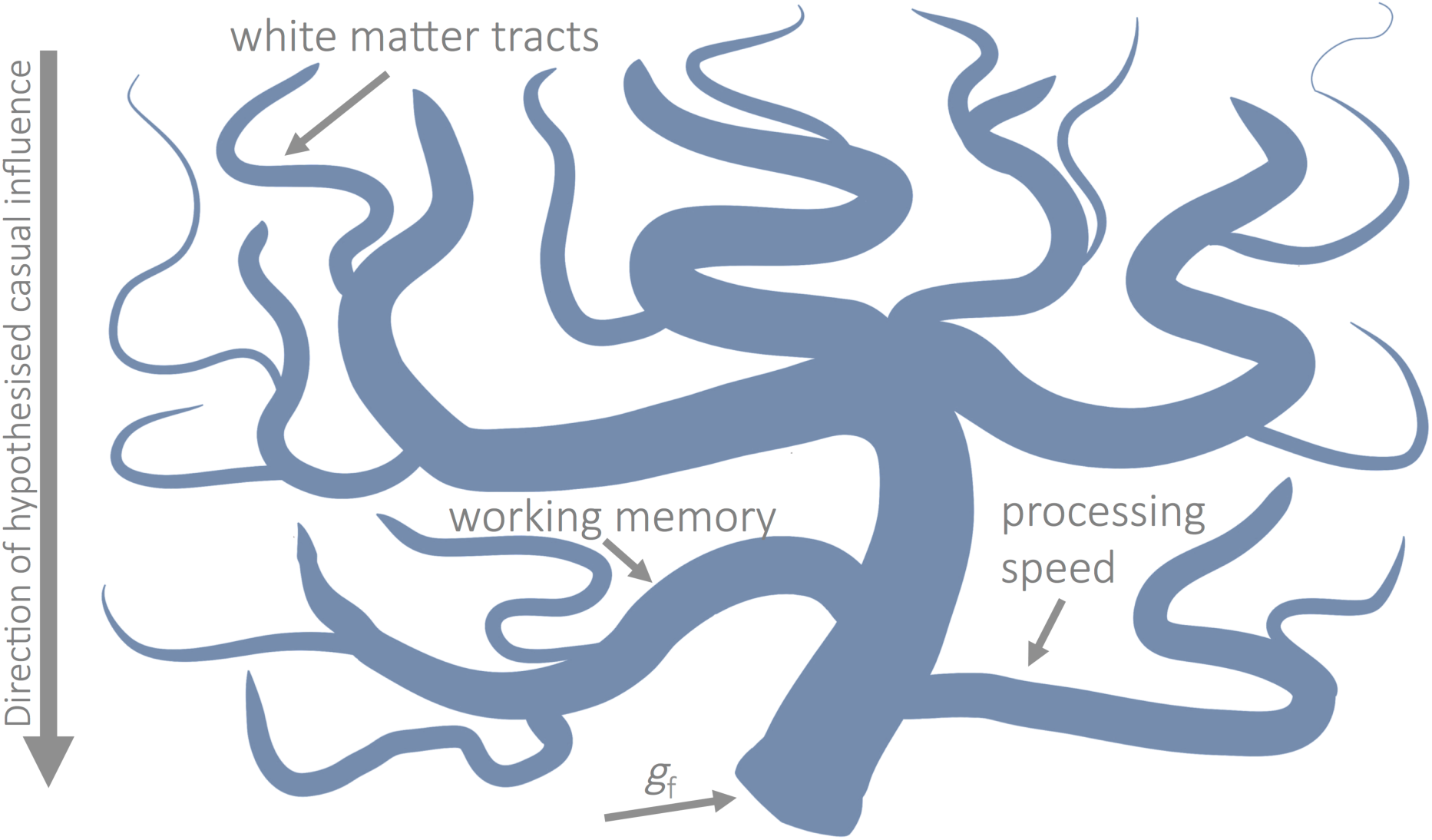
*The Watershed Model.* Schematic representation of the watershed model developed by (Cannon and Keller 2006) and adapted for the present study. Fluid ability is hypothesized to be the downstream product of working memory and processing speed, which are, in turn, the product of white matter contributions. Figure adapted from Kievit et al. (2016).

SEM, as a statistical technique, is uniquely suited to modeling the kinds of complex multivariate brain-behavior associations posited by the watershed model (Kievit et al. 2011; Kline 2015). SEM combines factor analysis and path analysis (a variant of regression analysis). It can model abstract cognitive constructs like *g*_f_, by estimating latent variables from observed task scores (i.e. manifest variables). This feature of SEM allowed us to model *g*_f_, working memory, and processing speed in two independent samples, and thereby provided a direct test of the generalizability of our findings. Second, SEM can test the simultaneous relations between multiple cognitive and neural variables, allowing us to address the relative contributions of different white matter tracts and different cognitive endophenotypes to *g*_f_. Finally, using SEM Trees (Brandmaier et al. 2013), a novel, decision-tree-based extension of SEM, we investigated whether the associations in the watershed model change with age.

Based on the watershed model we made the following preregistered predictions (http://aspredicted.org/blind.php?x=u5pf6z):

1. Working memory, *g*_f_ and processing speed are separable constructs.
2. Individual differences in *g*_f_ are predicted by working memory and processing speed.
3. White matter microstructure is a multi-dimensional construct.
4. There is a hierarchical relationship between white matter microstructure, cognitive endophenotypes (working memory and processing speed) and *g*_f_, such that white matter contributes to working memory and processing speed, which, in turn contribute to *g*_f_.
5. The contribution of working memory and processing speed to *g*_f_ changes with age.

## Materials and Methods

### Samples

We analysed data from the CALM and NKI-RS sample, as described in detail by (Holmes et al. 2018) and (Nooner et al. 2012) respectively. See also Simpson-Kent et al. (2019). We had also preregistered to analyse data from the ABCD cohort (Volkow et al. 2018). The latter cohort contains only data for 9 - and 10 - year olds at present, however, which limits comparability to CALM and NKI-RS, and makes it unsuitable for investigations of developmental differences. We therefore opted to not analyse ABCD data here and instead recommend a replication of the analyses presented here in ABCD once longitudinal data is available. The CALM sample consists of children and adolescents referred by health and educational professionals as having difficulties in attention, learning and/or memory. The NKI-RS is a community-ascertained, lifespan sample, and representative of the general population of Rockland, New York, and the United States as a whole, in terms of ethnicity, socioeconomic status etc. For NKI-RS, we included data for participants under the age of 18 only to match the age range of CALM and excluded data that were completed more than half a year after enrolment. The latter criterion was implemented to ensure that age at assessment did not differ substantively between cognitive measures. The final samples included 551 participants from CALM (30.85% female, aged 5.17 - 17.92 years, *N*_Neuroimaging_ = 165) and 335 participants from NKI-RS (43.48% female, aged 6.06 - 17.92 years, *N*_Neuroimaging_ = 67). See Table 1 for prevalence of relevant disorders and learning difficulties in the samples.

**Table 1.** Prevalence of Relevant Disorders and Learning Difficulties in the CALM and NKI-RS cohorts

| Variable | Percentage<br>CALM | Percentage<br>NKI-RS |
| --- | --- | --- |
| ADHD | 31.94 | 17.01 |
| Dyslexia | 5.81 | 5.67 |
| Autism | 6.72 | 0.60 |
| Mood disorder | 0.54 | 0.90 |
| Anxiety disorder | 2.36 | 18.21 |
| Medicated<sup>1</sup> | 10.53 | 17.01 |
| Speech/language problems | 38.11 | 19.40 |

*Note.* <sup>1</sup> unspecified medication for NKI-RS, ADHD-medication for CALM

### Cognitive Tasks

We included cognitive tasks measuring the domains of *g*_f_, working memory or processing speed for CALM and NKI-RS. See Table 2 for the complete list of tasks used, and the Supplementary Methods for task descriptions. Supplementary Figure 1 and 2 show raw scores on all tasks. The tasks modelled here were preregistered for CALM but not NKI-RS.

**Table 2.**
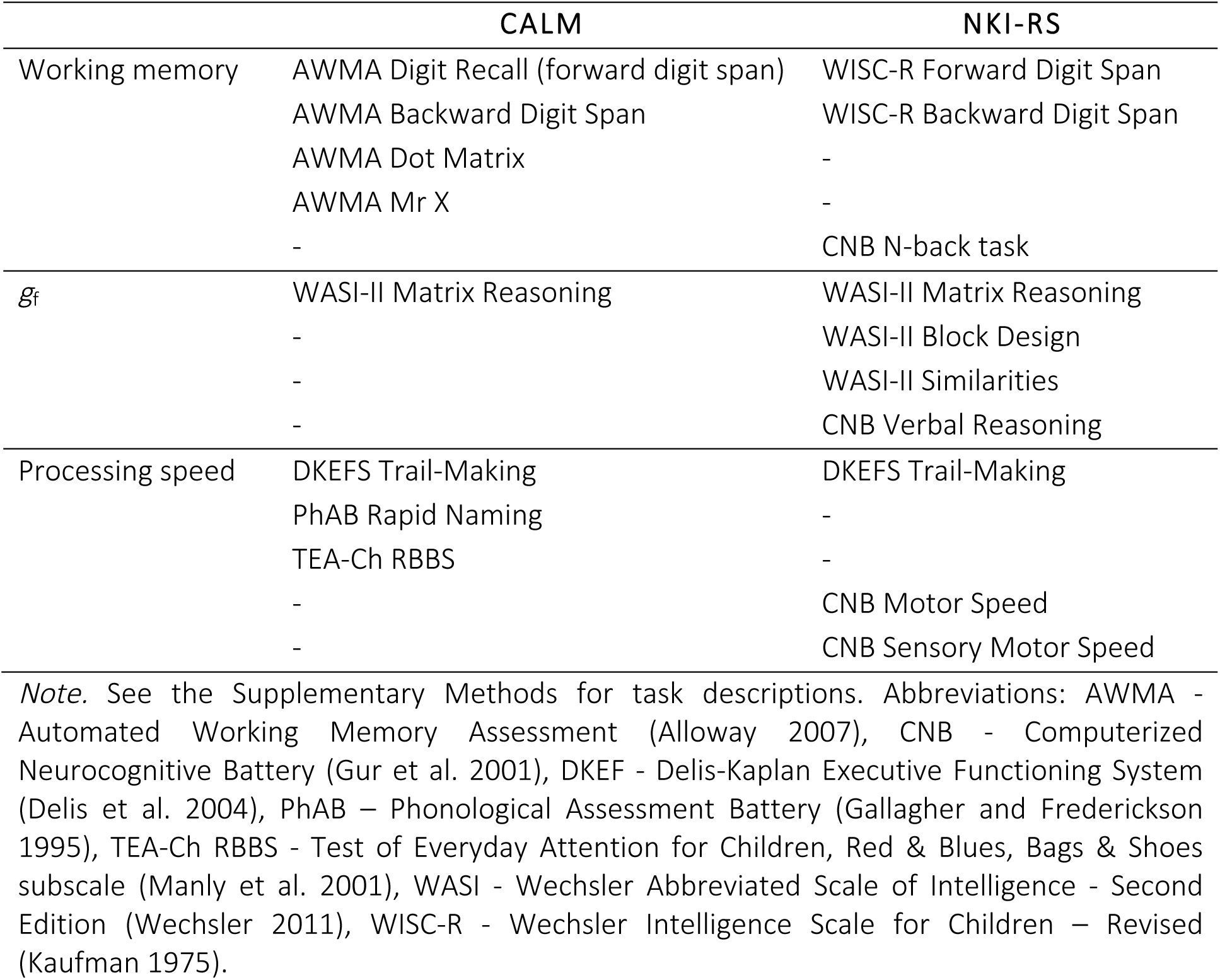
Cognitive Tasks Modelled

|  | CALM | NKI-RS |
| --- | --- | --- |
| Working memory | AWMA Digit Recall (forward digit span)<br>AWMA Backward Digit Span<br>AWMA Dot Matrix<br>AWMA Mr X<br>- | WISC-R Forward Digit Span<br>WISC-R Backward Digit Span<br>-<br>-<br>CNB N-back task |
| $g_f$ | WASI-II Matrix Reasoning<br>-<br>-<br>- | WASI-II Matrix Reasoning<br>WASI-II Block Design<br>WASI-II Similarities<br>CNB Verbal Reasoning |
| Processing speed | DKEFS Trail-Making<br>PhAB Rapid Naming<br>TEA-Ch RBBS<br>-<br>- | DKEFS Trail-Making<br>-<br>-<br>CNB Motor Speed<br>CNB Sensory Motor Speed |

*Note.* See the Supplementary Methods for task descriptions. Abbreviations: AWMA - Automated Working Memory Assessment (Alloway 2007), CNB - Computerized Neurocognitive Battery (Gur et al. 2001), DKEF - Delis-Kaplan Executive Functioning System (Delis et al. 2004), PhAB – Phonological Assessment Battery (Gallagher and Frederickson 1995), TEA-Ch RBBS - Test of Everyday Attention for Children, Red & Blues, Bags & Shoes subscale (Manly et al. 2001), WASI - Wechsler Abbreviated Scale of Intelligence - Second Edition (Wechsler 2011), WISC-R - Wechsler Intelligence Scale for Children – Revised (Kaufman 1975).

### White Matter Microstructure

We modelled mean FA for all ten tracts of the Johns Hopkins University (JHU) white matter tractography atlas (Hua et al. 2008) averaged over the hemispheres (Figure 2). See Supplementary Methods for details of the MRI acquisition and processing and Supplementary Figure 3 and 4 for raw FA values in all tracts.

**Figure 2.**
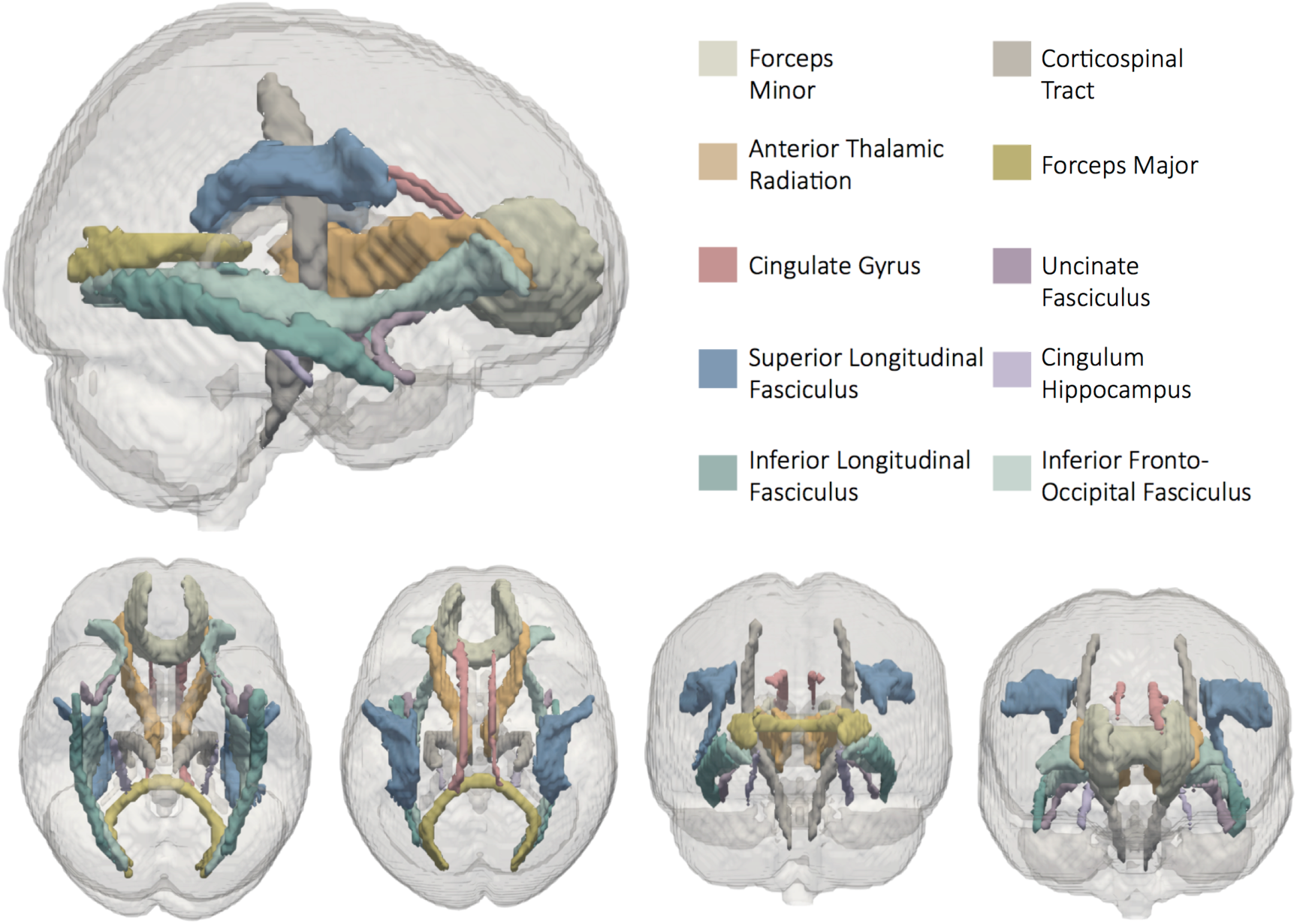
White Matter Tracts Modelled in the Analyses.

### Analysis Methods and Structural Equation Modelling

Covariance matrices and scripts replicating key analyses can be obtained from: https://github.com/df1234/gf_development. Supplementary Figure 5 and 6 show correlation matrices of all tasks and white matter tracts modelled. We modelled raw scores for *g*_f_ and working memory tasks, as preregistered. Raw scores on processing speed tasks were transformed. This step was not preregistered, but found necessary to achieve model convergence to ensure interpretability of scores. First, we inverted response time scores (using the formula y = 1/x) to obtain more intuitive measures of ‘speed’ for all but the CNB Motor Speed task, for which raw scores were already a measure of speed. Afterwards, we applied a log-transformation to reaction time tasks to increase normality and aid estimation. For the CNB Motor Speed task only, we additionally removed values ± 2 *SD* of the mean (*N* = 6) because the presence of these outliers had caused convergence problems.

We modelled the associations between cognition and white matter microstructure using SEM in R (R core team 2015) using the package lavaan (Rosseel 2012). All models were fit using maximum likelihood estimation with robust Huber-White standard errors and a scaled test statistic. Missing data was addressed using full information maximum likelihood estimation.

**Table 3.**
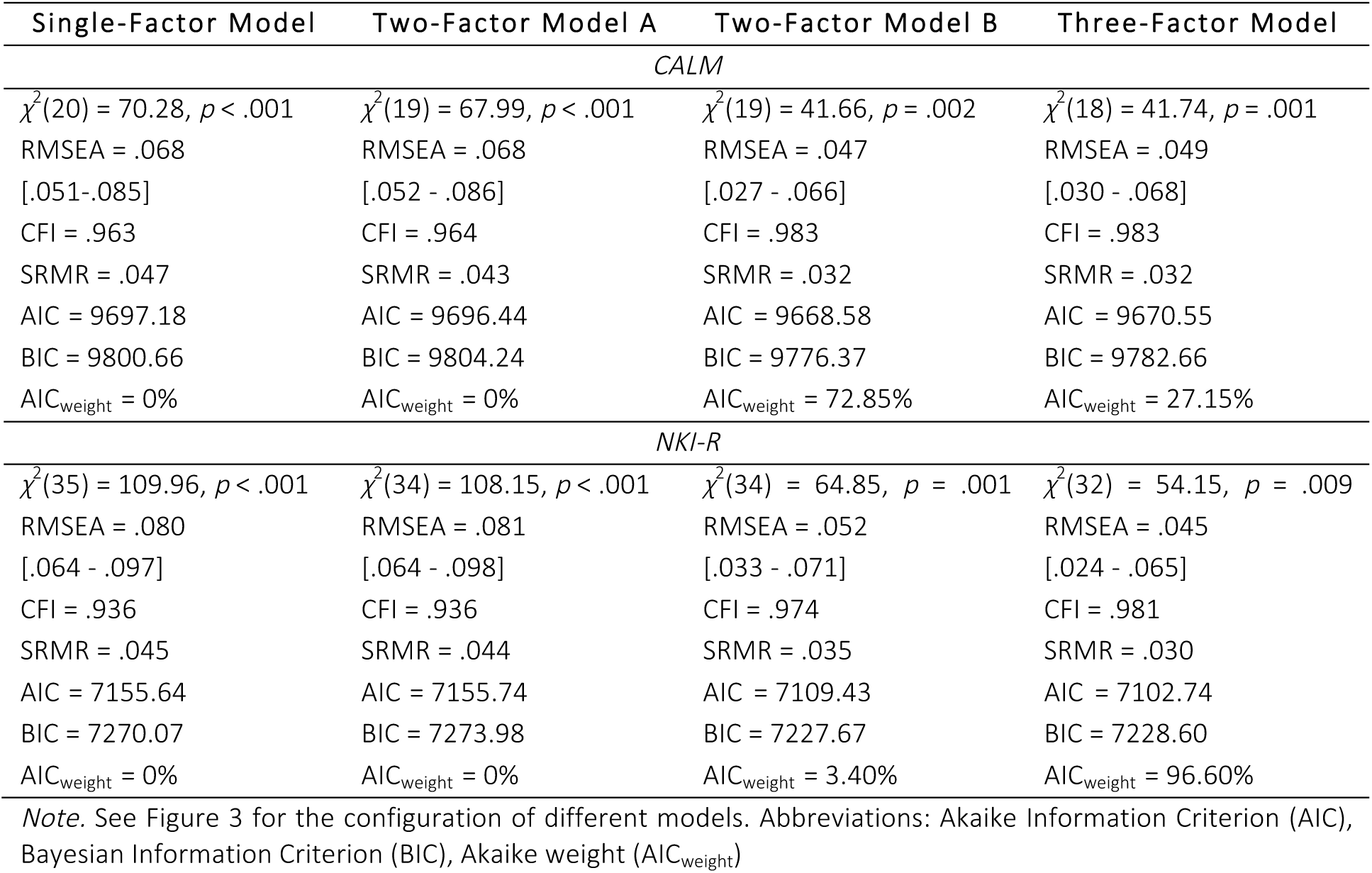
Model Fit of Competing Measurement Models

We used SEM Trees to investigate whether the associations among cognitive and neural measures differed with age. SEM Trees use decision tree methods to hierarchically split a dataset into subgroups if parameter estimates differ significantly based on a covariate of interest - in this case age (Brandmaier et al. 2013). We first ran the watershed model in OpenMx (Boker et al. 2011) and then passed this model object to semtree to compute the SEM Trees. We ran one SEM Tree for each parameter of interest (e.g. the covariance between working memory and processing speed). All other parameters in each semtree object were set to be invariant across groups to ensure that splits were specific to the parameter of interest. We used a 10 - fold cross-validation estimation method as recommend by (Brandmaier et al. 2013). For the path from the cingulate to working memory only we used 5 - fold cross-validation because the model did not converge using 10 - fold cross-validation. Minimum sample size in age group was set to *N* = 50 to ensure reliable estimation of standard errors. Note that this choice effectively limited search space for potential splits to ages 6.58 - 12.42 years for CALM and 8.08 - 15.49 years for NKI-RS.

## Results

To evaluate the hypotheses generated by the watershed model, we built up the watershed model in steps and carried our comprehensive tests of model fit at each step. First, we assessed the overall fit of our models to the data using the chi-square test, root mean square error of approximation (RMSEA), comparative fit index (CFI) and standardized root mean square residual (SRMR). Good absolute fit was defined as RMSEA < 0.05, CFI > 0.97 and SRMR < 0.05; acceptable fit as RMSEA = 0.08 - 0.05, CFI = 0. 95 - 0.97, SRMR = 0.05 - 0.10 (Schermelleh-Engel et al. 2003). Second, we assessed specific predictions from our models by comparing them to alternative models. Comparative model fit for nested models was assessed using the chi-square difference test. Non-nested models were compared using the Akaike (AIC) weights, which indicates the probability of a model being the data-generating model compared to all other models tested (Wagenmakers and Farrell 2004). Lastly, we evaluated the significance and strength of relationships between specific variables in our models by inspecting the Wald test for individual parameters, noting the joint *R*^2^ where relevant and reporting standardized parameter estimates. Absolute standardized parameter estimates above 0.10 were defined as small effects, 0.20 as typical and 0.30 as large (Gignac and Szodorai 2016).

### The Measurement Model of Cognition

To examine the neurocognitive architecture of *g*_f_, we started by modelling the cognitive components of the watershed model: *g*_f_, working memory and processing speed. Specifically, we fit a three-factor model of cognition (Figure 3) and compared it to alternative measurement models. This approach allowed us to test Hypothesis 1: namely that *g*_f_, working memory and processing speed form three separable, albeit likely correlated cognitive factors.

**Figure 3.**
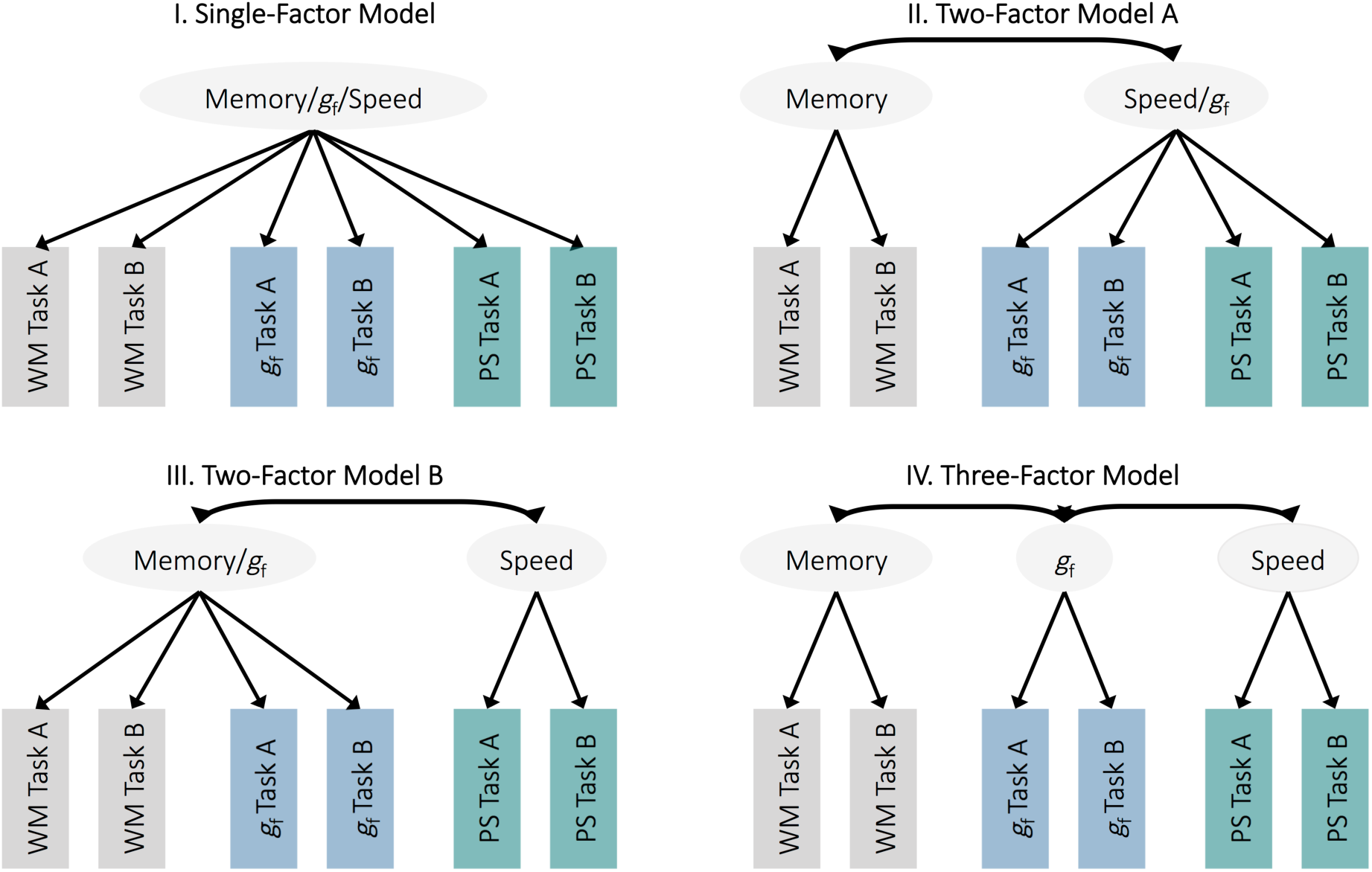
*Different Measurement Models of Cognition.* Abbreviations: WM: working memory, PS: processing speed

The Three-Factor Model (Figure 3) showed excellent *absolute fit* for both the CALM and NKI-RS sample (Table 3), indicating that overall, the data was compatible with a model of *g*_f_, working memory and processing speed as three separate factors.

The Three-Factor Model also showed very good *comparative fit* for NKI-RS, with a 96.60% probability of being the data-generating model compared to all alternative models tested, as indicated by its AIC weight (Figure 3). The evidence was more mixed for CALM, for which the Three-Factor Model showed a 27.15% probability of being the data-generating model, while Two-Factor Model B (Figure 3, treating working memory and *g*_f_ as a unitary factor) showed a 72.85% probability of being the data-generating model, highlighting a close relationship between *g*_f_ and working memory for this sample. The Single-Factor Model and Two-Factor Model A (Figure 3, treating speed and *g*_f_ as a unitary factor) showed a very low (approximately 0%) probability of being the data-generating model, indicating that speed and *g*_f_ were clearly separable in both samples.

Overall, these result provide mixed evidence for Hypothesis 1: Even though working memory, processing speed and *g*_f_ were highly correlated in both samples (Table 4), processing speed formed a clearly separable factor from working memory and *g*_f_ in both samples. Working and *g*_f_, however, were clearly separable only in NKI-RS, but not CALM, suggesting greater similarity between *g*_f_ and working memory in the CALM sample. To facilitate comparison across samples and in accordance with our preregistered analysis plan we nonetheless used the three-factor measurement model (Table 4, Supplementary Table 1) in all subsequent analyses.

**Table 4.** Covariance between Cognitive Measures in the Three-Factor Model

| Sample | Path | Standardized Estimate |
| --- | --- | --- |
| CALM | $g_f \leftrightarrow \text{memory}$ | 0.71, $z = 28.42$, $p < .001$ |
| | $g_f \leftrightarrow \text{speed}$ | 0.55, $z = 12.20$, $p < .001$ |
| | memory ↔ speed | 0.79, $z = 19.35$, $p < .001$ |
| NKI-RS | $g_f \leftrightarrow \text{memory}$ | 0.91, $z = 19.51$, $p < .001$ |
| | $g_f \leftrightarrow \text{speed}$ | 0.81, $z = 24.73$, $p < .001$ |
| | memory ↔ speed | 0.87, $z = 17.43$, $p < .001$ |

*Note.* See Supplementary Table 1 for factor loadings.

### The Relationship between Working Memory, Processing Speed and *g*_f_

We next examined the relationships between working memory, processing speed and *g*_f_ in more detail. Specifically, we fit a SEM including regression paths between working memory and *g*_f,_ as well as speed and *g*_f,_ to test Hypothesis 2, that working memory and processing speed each predict individual differences in *g*_f_. We found that this model showed good absolute fit for both samples (CALM: *χ*^2^(18) = 41.74, *p* = .001; RMSEA = .049 [.030 - .068]; CFI = .983; SRMR = .032, NKI-RS: *χ*^2^(32) = 54.15, *p* = .009; RMSEA = .045 [.024 - .065]; CFI = .981; SRMR = .030), indicating that, overall, the data was compatible with our model.

To further scrutinize the relationship between g_f_, working memory and speed, we compared our freely-estimated model to a set of alternative models with different constraints imposed upon the regression paths. First, to test whether working memory and speed each made different contributions, we tested an alternative model in which the paths from processing speed and working memory to *g*_f_ were constrained to be equal. In CALM (Δ*χ*^2^(1) = 15.53, *p* < .001), but not NKI-RS (Δ*χ*^2^(1) = 3.25, *p* = .072), the freely-estimated model fit better than the equality-constrained model, indicating that working memory and speed each made different contributions in CALM but not NKI-RS. Next, we tested whether the freely estimated model fit better than a model in which the path between *g*_f_ and working memory was constrained to zero. We found that that the freely estimated model fit better for both samples (CALM: Δ*χ*^2^(1) = 20.77, *p* < .001; NKI-RS: Δ*χ*^2^(1) = 12.97, *p* < .001). In line with our hypothesis, this result indicates that working memory makes a significant incremental contribution to *g*_f_. Finally, we tested a model in which the path between *g*_f_ and processing speed was constrained to zero. This model showed no difference in fit to the freely estimated model for CALM (Δ*χ*^2^(1) = 0.02, *p* = .875) or NKI-RS (Δ*χ*^2^(1) = 0.04, *p* = .849). Contrary to our hypothesis, this indicates that there was no clear incremental contribution of processing speed to *g*_f_.

Finally, we inspected standardized path estimates of the freely estimated model to assess the effect seizes of working memory and processing speed. Parameter estimates showed that working memory showed a greater effect on *g*_f_ than processing speed, particularly in CALM (Table 5) even though raw correlations between *g*_f_ and speed were high in both samples (Table 4).

**Table 5.** Regression Path Estimates.

| Sample | Path | Standardized Estimate |
| --- | --- | --- |
| CALM | speed -> $g_f$ | -0.01, $z = -0.16, p = .876$ |
| | memory -> $g_f$ | 0.72, $z = 7.65, p < .001$ |
| NKI-RS | speed -> $g_f$ | 0.06, $z = 0.21, p = .208$ |
| | memory -> $g_f$ | 0.86, $z = 1.81, p = .070$ |

Overall these results provide mixed evidence for Hypothesis 2: There was good evidence that working memory and speed made a significant joint contribution to *g*_f_, and that working memory made an incremental contribution to *g*_f_ in CALM. Contrary to our hypothesis, and the watershed model, however, processing speed showed no significant incremental contribution to *g*_f,_ above and beyond working memory. We explore likely explanations for this finding in the Discussion.

### The Measurement Model of White Matter

We next examined the measurement model of white matter to test Hypothesis 3, namely that white matter microstructure is a multi-dimensional construct. Specifically, we tested whether white matter microstructure could be adequately captured by a single factor by examining absolute model fit. As expected, the single-factor model of white matter microstructure did not fit the data well (CALM: *χ*^2^(35) = 124.63, *p* < .001; RMSEA = .125 [.103 - .147]; CFI = .933; SRMR = .039; NKI-RS: *χ*^2^(35) = 132.33, *p* < .001; RMSEA = .204 [.167 - .242]; CFI = .885; SRMR = .023). This indicates that white matter microstructure could not be reduced to a single ‘global FA’ dimension in our samples, in line with (Lövdén et al. 2013; Kievit, Davis, Griffiths, Correia, CamCAN, et al. 2016) and supporting Hypothesis 3. We therefore modelled each of the ten white matter tracts separately in all subsequent models.

### The Watershed Model: Relationships between Cognition and White Matter

Next, we fit the full watershed model including white matter, working memory, processing speed and *g*_f_. Following our general analysis procedure, we investigated overall model fit, alternative models and individual path estimates to gain a comprehensive understanding of the relationships in the watershed model and to test Hypothesis 4 - that white matter contributes to working memory capacity and processing speed, which, in turn, contribute to *g*_f_.

We found largely converging results across samples. The watershed model showed good absolute fit in CALM (*χ*^2^(78) = 107.78, *p* = .014; RMSEA = .026 [.012 - .038]; CFI = .981; SRMR = .043) and acceptable fit in NKI-RS (*χ*^2^(112) = 219.22, *p* < .001; RMSEA = .053 [.043 - .064]; CFI = .928; SRMR = .088). White matter explained large amounts of variance in working memory (*R*^2^_CALM_ = 32.3%; *R*^2^_NKI-RS_ = 46.1%) and processing speed (*R*^2^_CALM_ = 38.2%; *R*^2^_NKI-RS_ = 54.4%), which, in turn, explained even more variance in *g*_f_ (*R*^2^_CALM_ = 51.2%; *R*^2^_NKI-RS_ = 78.3%). In line with Hypothesis 4, this indicates that the watershed model fit the data overall.

Comparing the freely estimated watershed model to alternative, constrained, models showed that white matter contributed significantly to memory and processing speed. Specifically, a model in which paths from white matter to processing speed were constrained to zero fit worse than the freely-estimated model (CALM: Δ*χ*^2^(10) = 50.26, *p* < .001; NKI-RS: Δ*χ*^2^(10) = 27.19, *p* = .002), as did a model in which paths from white matter to working memory were constrained to zero (CALM: Δ*χ*^2^(10) = 52.26, p < .001; NKI-RS: Δ*χ*^2^(10) = 25.85, *p* = .004). As hypothesised, white matter therefore contributed to both processing speed and working memory.

We next inspected that relationship between individual white matter tracts and working memory and speed in more detail. A model in which paths from white matter to working memory and speed were constrained to be equal, fit worse than the freely-estimated watershed model for CALM (Δ*χ*^2^(18) = 47.76, *p* < .001) and NKI-RS (Δ*χ*^2^(18) = 30.42, *p* = .034), indicating that the role of white matter microstructure in supporting working memory and processing speed differed across tracts. This supports the notion that there is a many-to-one mapping between white matter and cognition - a core tenet of the watershed model.

Investigating individual standardised parameter estimates of the different white matter tracts showed that for CALM, only the anterior thalamic radiation contributed significantly to processing speed, whereas the superior longitudinal fasciculus, forceps major and cingulum were significantly, independently and positively related to working memory (Figure 4). For NKI-RS, the superior longitudinal fasciculus was significantly and positively related to processing speed and working memory (Figure 5). Two tracts showed an unexpected, strongly negative (< −1), relationship: the forceps minor for CALM and the inferior fronto-occipital fasciculus for NKI-RS. We found that these negative estimates occurred only when all other brain to cognition pathways were also estimated: When estimated on their own, path estimates were positive (forceps minor to working memory: standardized estimate = 0.36, *z* = 4.05, *p* < .001; inferior fronto-occipital fasciculus to working memory: standardized estimate = 0.14, *z* = 0.859, *p* = .390; inferior fronto-occipital fasciculus to processing speed: standardized estimate = 0.26, *z* = 1.41, *p* = .158). This sign-flip suggests that the negative pathways were potentially due to modelling several, highly-correlated paths at the same time (Jöreskog 1999). Overall, these results further support the watershed prediction that multiple white matter tracts map onto working memory and processing speed.

**Figure 4.**
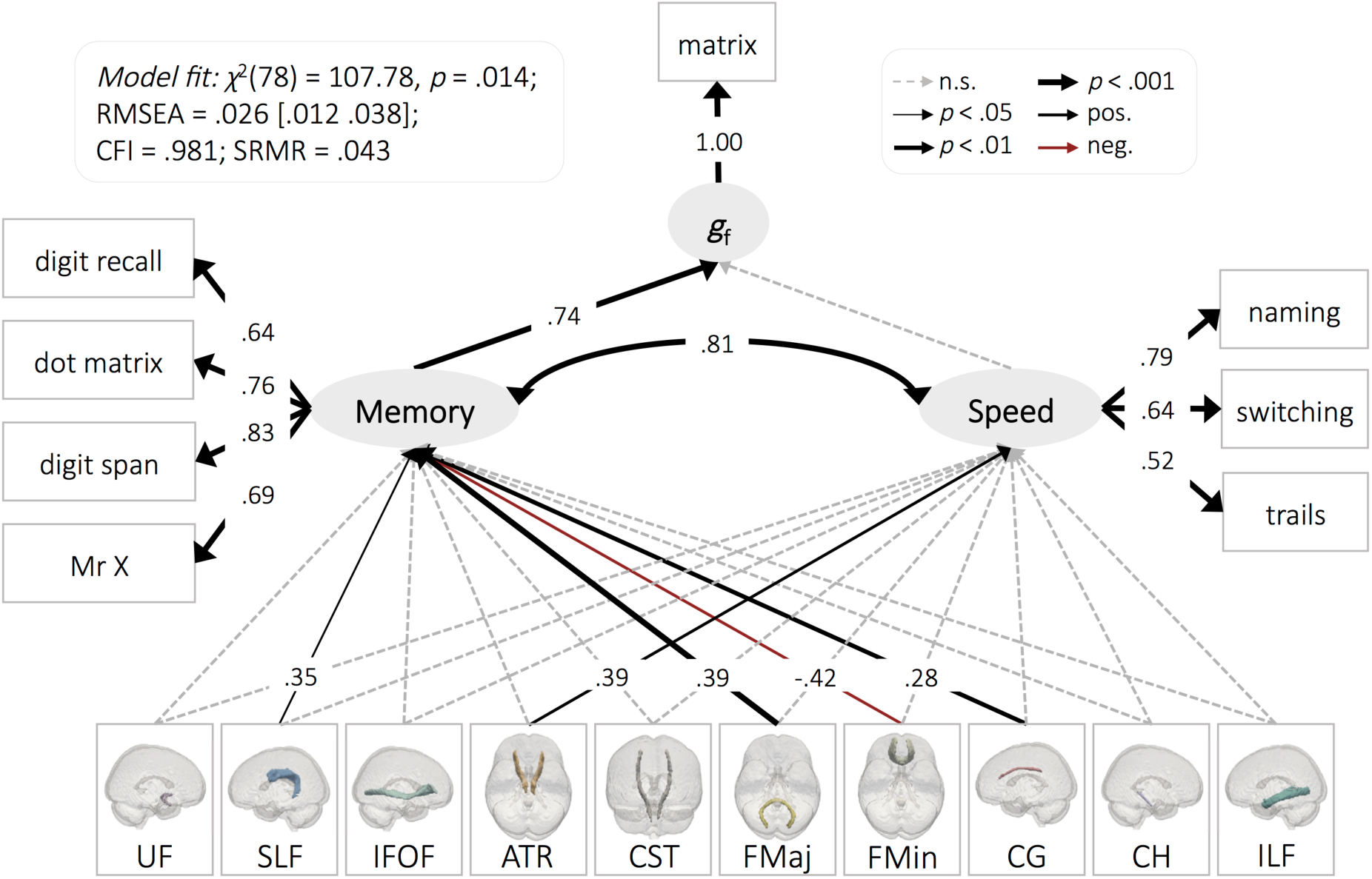
*The Watershed Model in CALM.* See Supplementary Table 2 for regression estimates. Residual covariances between white matter tracts were allowed but are not shown for simplicity. Abbreviations: uncinate fasciculus (UF), superior longitudinal fasciculus (SLF), inferior fronto-occipital fasciculus (IFOF), anterior thalamic radiations (ATR), cerebrospinal tract (CST), forceps major (FMaj), forceps minor (FMin), dorsal cingulate gyrus (CG), ventral cingulate gyrus (CH), inferior longitudinal fasciculus (ILF).

**Figure 5.**
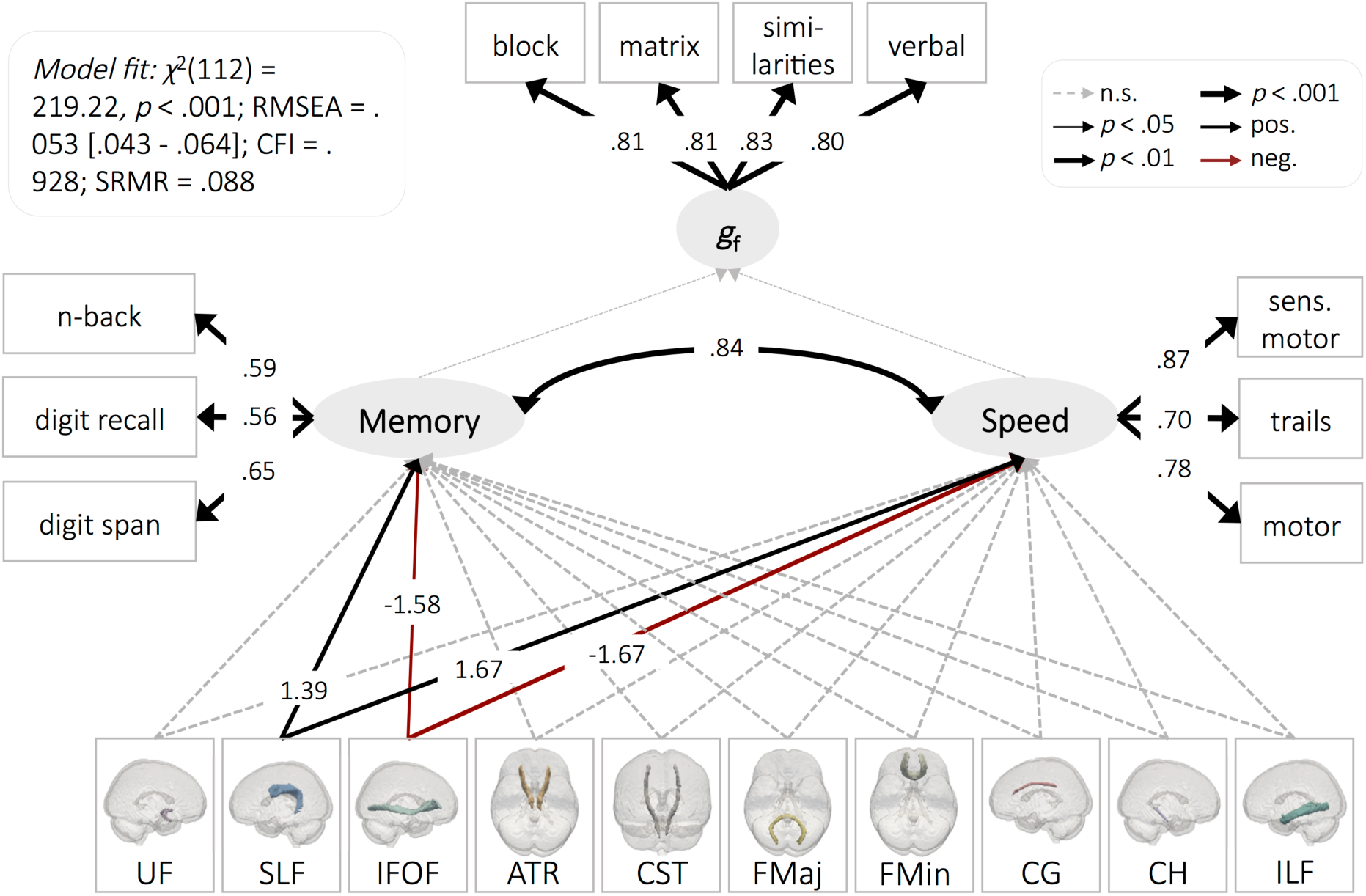
*The Watershed Model in NKI-RS.* See Supplementary Table 3 for regression estimates. Residual covariances between white matter tracts were allowed but are not shown for simplicity.

Finally, we probed the watershed model in more detail by testing a set of alternative expressions of the watershed model still compatible with the core tenants of the watershed model – as well as a set of alterative models incompatible with the watershed model. We compared all alternatives (see Figure 6 for graphical representations) to the original watershed model by inspecting their relative probability of being the data-generating model as indicated by their AIC weights (Wagenmakers and Farrell 2004). We found that the original watershed model showed a very high probability (98.58%) of being the data-generating model for CALM but only a 0.10% probability for NKI-RS. For NKI-RS, a different expression of the watershed model, such that *g*_f_ was regressed on working memory, which was regressed on processing speed, which was then regressed on white matter (Alternative A, Figure 6) showed a 95.04% probability of being the data-generating model. This model only showed a 0.37% probability for CALM. Another expression of the watershed model, in which all tasks were modelled separately as manifest, rather than latent, variables (Alternative B, Figure 6), showed no advantage over the watershed model for CALM (0.00% probability) or NKI-RS (0.00% probability). We next tested two alternative models incompatible with the tenants of the watershed model. We found that a model in which the hierarchy between cognitive endophenotypes and *g*_f_ was inverted (Alternative C, Figure 6) showed comparatively low probability of being the data-generating model for both CALM (0.00%) and NKI-RS (2.86%). Similarly, a model in which *g*_f_ was directly regressed on white matter, working memory and processing speed (Alternative D, Figure 6), showed no clear advantage over the watershed model for CALM (1.05% probability) or NKI-RS (0.00% probability). Overall these model comparisons highlight that while the watershed model fit the data for both samples and had large explanatory power (as indicated by *R*^2^s), the precise configuration of the watershed model may differ somewhat between cohorts.

**Figure 6.**
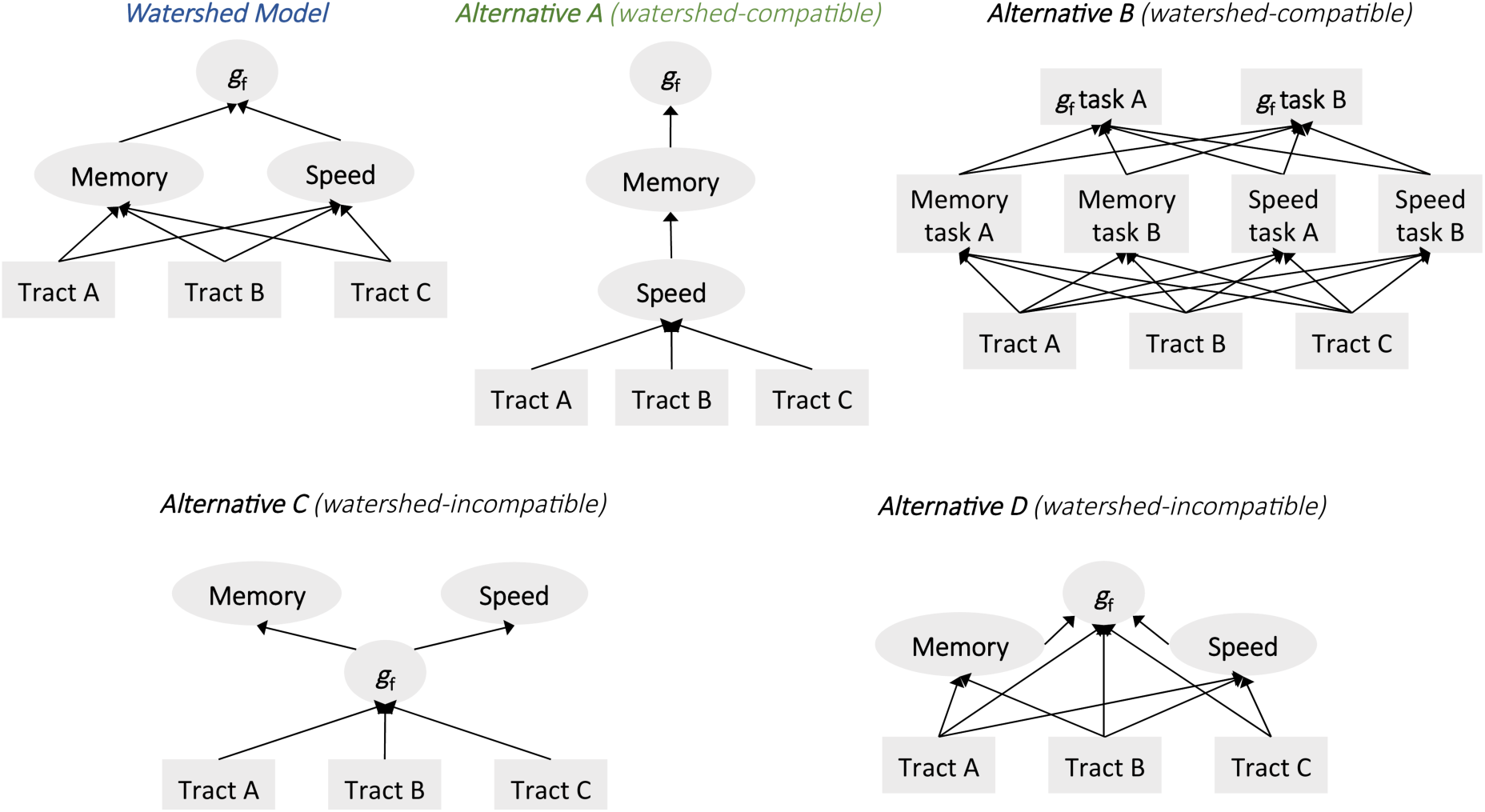
*Configuration of Alternative Models.* Alternatives A and B are watershed-compatible, while C and D are watershed-incompatible. The best-fitting model for CALM is highlighted in blue; the best-fitting model for NKI-RS is highlighted in green. Regression paths only are shown for simplicity. Square shapes denote manifest variables and oval shapes latent variables.

In summary, we found that the watershed model performed well overall for both cohorts. As hypothesised, white matter contributed to working memory and processing speed, which, in turn, contributed to *g*_f_, and explained large amounts of variance therein. Also as predicted by the watershed model, there was a many-to-one mapping between white matter tracts and cognition. The exact configuration of the watershed model, however, may differ slightly between cohorts. These differences may be a function of cohort differences in sample size, average levels of cognitive ability and/or the specific tasks used – a topic we will return to in the Discussion.

### Testing for potential confounds

We carried out a series of supplementary and non-preregistered analyses to examine whether possible confounders influenced our models. These analyses showed that our findings were robust to the inclusion of covariates such as scanner motion or socio-economic status. They were also robust across genders and participants taking or not taking medication. There were no differences in the structure of the model between participants with and without diagnosed disorders for CALM. Potential small differences cannot be ruled out for NKI-RS, likely due to the low number of diagnosed participants of *N* = 106 (Supplementary Material).

### Age-Related Differences in the Neurocognitive Architecture of *g*_f_

Finally, we tested Hypothesis 5 - that that the contribution of working memory and processing speed to *g*_f_ varied with age. We first inspected cross-sectional differences in *g*_f_, working memory and processing speed, and then used SEM trees to investigate potential age-differences in the relationships between these factors. In additional, non-preregistered, analyses we also used SEM Trees to investigate potential age-differences in the relationship between white matter and cognitive endophenotypes by inspecting paths that were significant in the watershed model (Figure 4 and 5).

SEM trees combine SEMs with decision tree methods, separating a dataset into subgroups (in this case age groups) if SEM parameter estimates of interest differ sufficiently (Brandmaier et al. 2013). SEM trees allowed us to investigate age as a potential moderator without imposing a-priori categorical age splits. We initially allowed for no more than two age groups. This yielded inconsistent results for CALM and NKI-RS (Supplementary Table 4). To test whether these inconsistencies were an artefact of allowing for only two groups, we repeated our analysis and allowed for up to four age groups. This analysis yielded consistent results between CALM and NKI-RS (Table 6). This pattern of results indicates that the initial parameters of our analysis caused us to miss relevant age differences.

**Table 6.** SEM Tree Results for the Watershed Model.

| Path | Est.<br>Before | Age<br>Split<br>1 | Est.<br>Betw. | Age<br>Split<br>2 | Est.<br>Betw. | Age<br>Split<br>3 | Est.<br>After |
| --- | --- | --- | --- | --- | --- | --- | --- |
| *CALM* |  |  |  |  |  |  |  |
| memory <→ speed | 0.85 | **8.46** | 0.97 | **9.46** | 0.74 | - | - |
| memory → $g_f$ | 0.83 | **9.38** | 0.42 | **10.04** | 1.14 | **10.88** | 0.94 |
| speed → $g_f$ | 0.04 | **6.88** | -0.19 | **11.21** | 0.17 | - | - |
| SLF → memory | 0.67 | **7.21** | 0.18 | **11.21** | 0.76 | - | - |
| FMaj → memory | 0.59 | **7.71** | 0.14 | **9.29** | 0.33 | **11.13** | 0.74 |
| CG → memory<sup>1</sup> | 0.64 | **6.96** | 0.09 | **11.04** | 0.70 | - | - |
| ATR → speed | 0.96 | **7.13** | 0.68 | **7.96** | 0.17 | **11.96** | 0.65 |
| *NKI-RS* |  |  |  |  |  |  |  |
| memory <→ speed | 0.90 | **9.82** | 0.48 | **14.72** | 1.11 | - | - |
| memory → $g_f$ | 1.10 | **8.59** | 0.59 | **12.67** | 1.03 | - | - |
| speed → $g_f$ | 0.53 | **8.59** | -0.12 | **12.96** | 0.52 | - | - |
| SLF → memory | 2.15 | **8.30** | 1.47 | **12.15** | 1.93 | - | - |
| SLF → speed | 3.12 | **8.63** | 1.83 | **15.09** | 2.31 | - | - |

*Note.* The table shows differences in parameter estimates for paths of interest (as shown in Figure 4 and 5) depending on participants' age in years. Our analyses allowed for a maximum of three age splits (and thus four age groups). An absence of a third age split (denoted by '-' in the table), indicates that the SEM tree split only twice, suggesting no further changes in parameter strength after the second split. See Supplementary Figure 7 for a graphical representation of these results.

As shown in Figure 7, *g*_f_, working memory and processing speed factor scores increased with age for all three cognitive phenotypes. In line with our hypothesis, SEM trees showed that there were pronounced age-related differences in brain-behaviour in childhood and adolescence (Table 6). For both samples and all but one path, there was an initially strong relationship between components of the watershed model, then a dip around ages 7 - 9 years for CALM and age 8 for NKI-RS, followed by an increase in path strength around ages 11 – 12 (see Supplementary Figure 7 for a graphical representation of these results). Speculatively, this pattern of results is consistent with an interpretation of a reorganization of neurocognitive faculties in late childhood, followed by a consolidation of neurocognitive pathways around the onset of adolescence (Johnson 2000, 2011).

**Figure 7.**
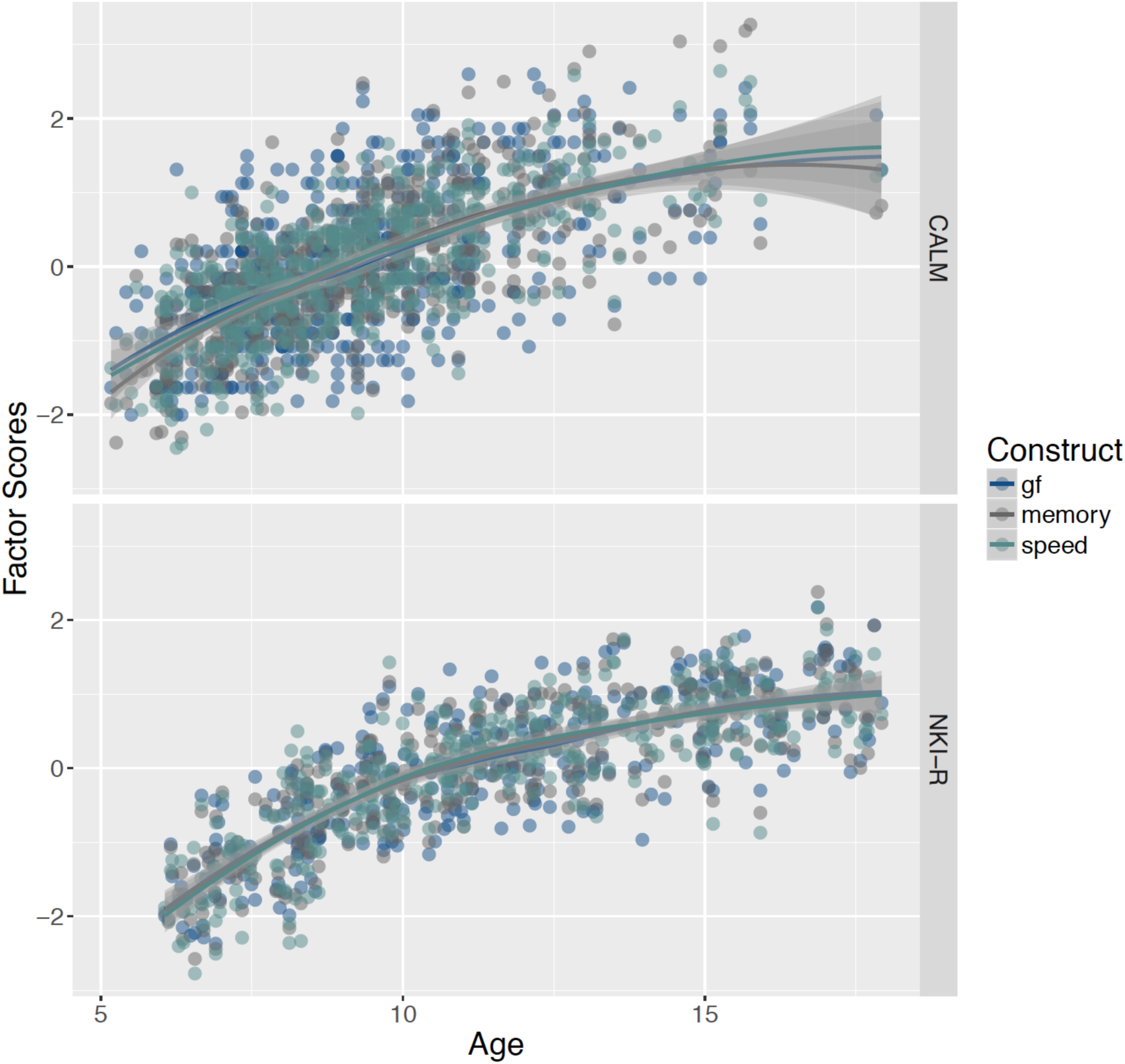
Cognitive Factor Scores by Age.

## Discussion

We here used multivariate statistical techniques to investigate the neurocognitive architecture of *g*_f_ in two large (*N*_CALM_ = 551, *N*_NKI-RS_ = 335) developmental cohorts and, for the first time, investigated how the neurocognitive architecture of *g*_f_ changes dynamically with age. We tested a preregistered watershed model of *g*_f_, which predicts a hierarchy of partially independent effects. As might be expected from a multi-cohort study, there were some differences between the community-ascertained cohort (NKI-RS) and the cohort of children and adolescents with learning difficulties (CALM) in specific path estimates. Overall however, we found strikingly convergent results across these two heterogeneous samples. The watershed model performed well for both CALM and NKI-RS: White matter contributed to working memory and processing speed, which, in turn, contributed to *g*_*f*_ and explained 51% of variance therein for the CALM sample and 78% of variance for NKI-RS. Models were robust across genders, participants taking or not taking medication and when controlling for socio-economic status and scanner motion. Investigations of age effects showed that the relationship between cognitive abilities and white matter dipped in strength around ages 7-12 years. Speculatively, this age-effect may reflect a reorganization of the neurocognitive architecture during pre-puberty and early puberty (Byrne et al. 2017). These findings have implications for understanding and targeting cognitive impairments in populations with learning difficulties.

The watershed model tested here consists of three levels: *g*_f_ forms the most down-stream point, with working memory and processing speed as intermediate tributaries, and white matter microstructural tracts as upstream sources. Previous studies suggested that matter microstructure is best characterised by a single, ‘global FA’ factor (Penke et al. 2010) while others have contended that association patterns among different white matter tracts are more complex (Lövdén et al. 2013; Kievit, Davis, Griffiths, Correia, CamCAN, et al. 2016). Here we found strong evidence for a multifactorial view of white matter tracts – for both samples, a unidimensional model of white matter fit poorly, and for CALM, multiple tracts also showed partially independent contributions to distal cognitive outcomes. This is in line with the watershed model. There were some differences between cohorts as to which tracts contributed most to working memory and processing speed: In line with previous research (Kievit, Davis, Griffiths, Correia, CamCAN, et al. 2016; MacPherson et al. 2017; Bathelt et al. 2018), we found that the anterior thalamic radiation was related to processing speed, as were the forceps major, forceps minor and the cingulum to working memory for CALM. However, these tracts were not significant for NKI-RS. These differences between samples may reflect differences in brain-behaviour mapping between more atypical and typical cohorts (Bathelt et al. 2018), or simply sampling variance across two independent cohorts where one cohort (NKI-RS) has a more modest number of participants with imaging data. Of note, however, the superior longitudinal fasciculus was consistently associated with working memory in both samples. For NKI-RS, the superior longitudinal fasciculus was also associated with processing speed. The superior longitudinal fasciculus is a large, bilateral association fibre connecting temporal, occipital, parietal and frontal regions (Kamali et al. 2014). It is therefore well-situated for supporting cognitive processes such as *g*_f_, which rely on integrative multiple-demand systems (Jung and Haier 2007; Fedorenko et al. 2013; Parlatini et al. 2017).

The cognitive levels of the watershed model highlighted a close relationship between working memory and *g*_f_. Previous studies had variably suggested that *g*_f_ and working memory (Kyllonen and Christal 1990; Fukuda et al. 2010), or *g*_f_ and processing speed (Kail and Salthouse 1994; Salthouse 1996; Coyle et al. 2011; Ferrer et al. 2013) may be most closely related. We found that all three cognitive factors were highly correlated for both samples. Nonetheless, processing speed formed a cognitive factor clearly separable from working memory and *g*_*f.*_ Working memory and *g*_f_, in turn, were separable in the community-ascertained NKI-RS but not in CALM, the cohort of children and adolescents with learning difficulties. This close relationship between *g*_f_ and working memory was also evident in other models of CALM where processing speed and working memory were used as joint predictors of *g*_*f*_: Contrary to our hypotheses, processing speed became non-significant after controlling for working memory here. There are several possible, and not mutually exclusive, explanations for this finding. First, and in line with previous work showing that time-constraints increase isomorphism of *g*_*f*_ and working memory (Chuderski 2013), even standard implementations of *g*_*f*_ tasks may place considerable time-pressure on struggling learners, whereby increasing *g*_*f*_ - working memory covariance. Second, a broader set of speed tasks (which might be captured by several latent variables for clerical speed tasks, choice reaction time tasks and variability indices) might show higher predictive power than the single latent variable for speed, which could be modelled here. Third, the watershed model might be configured somewhat differently for some populations, such that speed forms an intermittent level in the hierarchy between white matter and working memory (Alternative A, Figure 6). There was some evidence for this in NKI-RS, indicating that the hierarchy of the watershed model might be differentiated more in cohorts of older ages and/or higher ability levels. We note that all of these explanations would still be compatible with the notions of the watershed model, and remain to be teased apart by future research. For now, we suggest that our findings support the notion that mental information processing capacity, as measured by working memory, is a key determinant of *g*_f_ (Kyllonen and Christal 1990; Fukuda et al. 2010).

The associations in the watershed model differed between ages in a complex, non-monotonic fashion. Previous research had suggested either a decrease in covariance among cognitive domains with age (age differentiation) (Garrett 1946), an increase in covariance with age (age de-differentiation) (Blum and Holling 2017), or no changes with age (Tucker-Drob 2009; de Mooij et al. 2018). These investigations have traditionally focussed on relations between cognitive domains, however, not on relationships between brain and cognition - although see de Mooij et al. (2018). Possible linear and non-linear changes in brain-behaviour mapping with age have remained mostly unexplored (Tamnes et al. 2017). Using structural equation modelling trees, a novel decision-tree-based technique, we here found evidence of complex developmental differences that were consistent across samples and relationships in the watershed model: Initially strong path estimates showed a pronounced decrease in strength around ages 7 - 9 years, followed by a renewed increase in the strength, even surpassing initial levels, around ages 10 - 15.

There are at least two possible explanations for this developmental dip in brain-cognition relationships. First, there may be a true decrease in relationship strength during this time of life. Possibly, other cognitive skills such as verbal reasoning temporarily support *g*_f_, resulting in weaker relationships between *g*_f_ and working memory, for instance. Alternatively, the configuration of the watershed model may change temporarily during this time, which could also manifest in an apparently weaker covariance structure. In this case, the true relationship between *g*_f_, memory, speed and white matter may still be strong, just configured differently from the watershed model. Both explanations are compatible with the interactive specialization theory (Johnson 2000, 2011), which predicts as remapping of the relationships between brain substrates and cognitive abilities during development.

On a physiological level, this age effect may be driven by neuroendocrine changes during pre- and early puberty. Puberty is driven by a complex and only partially understood set of hormonal events including gonadarche and andrenarche (Sisk and Zehr 2005). Gonadarche begins with the secretion of gonadotropin-releasing hormone from the hypothalamus around ages 10-11 years and closely tracks the overt bodily changes of puberty (Dorn 2006). Andrenarche, beginning with the maturation of the andrenal gland, starts as early as six years of age, and is increasingly recognized as a complimentary driver of puberty and brain development (Byrne et al. 2017). It is possible that the hormonal changes of andrenarche and early gonadarche may lead to a level of neural reorganization, which may initially appear as weaker relationships in the watershed model. The sweeping bodily, social and cognitive changes happening in early adolescence may then drive a consolidation of the neurocognitive architecture of *g*_f_.

On a more general level, this age effect suggests the existence of potential non-linear changes in brain-behaviour mapping during childhood and adolescence and underlines the value of modern statistical approaches, such as SEM Trees, for the study of age-related differences. Nonetheless, it is worth noting that these findings, based on an inherently exploratory technique, will need to be replicated in future confirmatory studies with fine-grained data on puberty and larger sample sizes. The latter will also allow for detailed investigations of potential gender differences.

Testing our model in two different samples allowed us to address several critical questions: First, participants from both samples completed a small set of common and a larger set of different cognitive tasks. Therefore, the results obtained here are likely not task-specific, but rather can be expected to generalize to the domains of working memory, processing speed and *g*_*f*_ (Noack et al. 2014). For instance, we show that even though one cohort (NKI-RS) performed mainly clerical speed tasks and the other (CALM) mainly choice reaction time tasks, our finding that processing speed did not significantly contribute to *g*_f_ after controlling for working memory, was robust across these samples and tasks. Second, by comparing a community-ascertained sample (NKI-RS) and a sample of children and adolescents with learning difficulties (CALM), we demonstrated that the watershed model performed well for very different populations. We are not able to make general claims about potential differences between more typical and atypical populations, however. CALM and NKI-RS were collected in countries with somewhat different socio-economic conditions (United Kingdom and United States of America), the samples were of different sample sizes, and participants’ ages were distributed more evenly in one cohort (NKI-RS) than the other (CALM). While DTI images were processed with the same pipeline across sites, the scanner and MRI acquisition protocol were different. Although previous work suggests that FA is relatively robust measure in multi-site comparisons (Vollmar et al. 2010), we cannot rule out site differences as a potential confound. It will therefore be necessary to replicate these findings in large typical and atypical cohorts collected in the same setting.

Our study illustrates some of the advantages and challenges of preregistered secondary data analyses. We agree with others in the field that secondary data analysis need not be and should not be confounded with purely exploratory research (Mills and Tamnes 2014; Orben and Przybylski 2019; Scott and Kline 2019). Preregistrations, as well as dedicated multivariate methods such as SEM, can help to reduce the scope for analytic flexibility and increase scientific rigour when using secondary datasets, which are often rich and multivariate in nature. Preregistrations also do not preclude the use of exploratory methods (such as SEM trees used here) or the ability to ask exploratory questions (such as looking at age differences in the relationships between brain and cognition reported here) - they merely facilitate the distinction between exploratory and confirmatory research (Wagenmakers et al. 2012). There are, however, some unique challenges to preregistering secondary data analyses worth noting. First, information on the precise measures collected is not always easily available prior to data access, which can limit the level of detail in which an analysis can be preregistered. Second, data quality and the level of data-processing (the latter is particularly relevant for MRI data) is not always clear a priori (e.g. see Kievit et al. 2018), which can necessitate changes to analyses plans after data inspection. Third, convergence issues are fairly common when using complex multivariate methods such as SEM. We found it necessary to transform some of our speed variables, for instance, to achieve model convergence. Such post-hoc modifications, not guided by the palatability of the results, but rather by unforeseen, and sometimes unforeseeable, practical considerations, mean that preregistration can sometimes fall short of full compliance. Nevertheless, we believe that even imperfect preregistrations, alongside shared code, data and the transparent presentation of results, can help the reader distinguish between confirmatory and exploratory results, and adjust their level of confidence in conclusions accordingly. For guidance on maximizing transparency in preregistration of secondary data, see Weston et al. (2018).

A key limitation of our study is that our samples were cross-sectional, and not longitudinal. Although the relatively narrow age range makes large cohort effects unlikely, it may still be that there were differences in recruitment and selection that varied across the age range. Moreover, while we were able to investigate *individual differences* in *g*_f_, we could not assess *intra-individual changes* during childhood and adolescence. As such, the cross-sectional nature of our samples limits our ability to make inferences about developmental dynamics. Longitudinal data will also be necessary to scrutinize the causal flow of effects in the watershed model.

The findings from our study have implications understanding and targeting cognitive impairments in populations with learning difficulties. First, the close relationship between working memory and *g*_f_ found here and in other studies (Fukuda et al. 2010; Chuderski 2013), indicates that children and adolescents struggling with working memory are likely to also struggle in terms of complex reasoning tasks. Either reducing working memory load, decreasing time constraints, or training working memory and fluid ability capacity in such populations may therefore be promising lines of inquiry for intervention studies. It is worth highlighting, however, that cognitive training studies have so far shown little evidence of (far) transfer: Training abstract reasoning, a common measure of *g*_*f*_, has not resulted in robust increases in working memory (Knoll et al. 2016) and working memory training has not been shown to transfer to reasoning skills or school performance (Dunning et al. 2013; Schwaighofer et al. 2015). Similarly, transfer from processing speed to reasoning seems to be limited (Mackey et al. 2011). The results obtained here suggest that interventions may increase their chance of success by implementing programs of sufficient complexity to affect the entire neurocognitive architecture of effects (see also Kievit et al. 2016). The level of intensity required to produce sustained benefits may need to be as demanding and consistent as education itself, which shows robust effects in increasing general cognitive abilities over time (Ritchie and Tucker-Drob 2018). This work and work by others (Noack et al. 2014) also highlights the value of assessing, modeling, and potentially intervening on, multiple tasks, rather than relying on a single task to capture complex cognitive domains such as *g*_f_. Finally, the age-related differences in the relationships of the watershed model observed using SEM-trees suggest that some interventions may work best at particular developmental phases.

## Acknowledgements

D.F. and J.B. are supported by the UK Medical Research Council (MRC). I.L.S.-K. is supported by the Cambridge Trust. R.A.K. is supported by the Sir Henry Wellcome Trust (107392/Z/15/Z) and MRC Programme Grant (MC-A060-5PR60). The Centre for Attention Learning and Memory (CALM) research clinic is based at and supported by funding from the MRC Cognition and Brain Sciences Unit, University of Cambridge. The Principal Investigators are Joni Holmes (Head of CALM), Susan Gathercole (Chair of CALM Management Committee), Duncan Astle, Tom Manly and Rogier Kievit. Data collection is assisted by a team of researchers and PhD students at the CBU that includes Sarah Bishop, Annie Bryant, Sally Butterfield, Fanchea Daily, Laura Forde, Erin Hawkins, Sinead O’Brien, Cliodhna O’Leary, Joseph Rennie, and Mengya Zhang. The authors wish to thank the many professionals working in children’s services in the South-East and East of England for their support, and to the children and their families for giving up their time to visit the clinic. We would also like to thank all NKI-RS participants and researchers. We are grateful to Amber Ruigrok for helpful suggestions regarding pubertal development.

