## Supplementary Material for "A hierarchical watershed model of fluid intelligence in childhood and adolescence"

### Supplementary Methods

#### Cognitive Tasks.

*AWMA Digit Recall/ WISC-R Forward Digit Span.* In this verbal short-term memory span task, participants were asked to repeat sequences of single digit numbers presented in audio format.

*AWMA Backward Digit Span / WISC-R Backward Digit Span.* This task was similar to the digit recall task but participants were asked to recall the sequences of numbers in backward order.

*AWMA Dot Matrix.* In this visuo-spatial working memory span task, participants were presented with a sequence of red dots shown in a grid. They were then asked to recall the position of the dots at the end of each sequence.

*AWMA Mr X.* In this second visuo-spatial working memory span task, participants were shown two cartoon figures (Mr Xs), each carrying a ball in one hand. For each trial, the left Mr X figure stood upright while the right Mr X figure was rotated to different positions. Participants were then instructed to recall the location of the ball held by the right Mr X after it disappeared and whether he held the ball in the same hand as the left Mr X.

*CNB N-back task.* In this verbal working memory task, participants were shown letter sequences presented for 500 ms each and asked to respond by pressing a key according to the following rules: 1-Back – participants pressed whenever the letter presented was identical to the previous letter; and, 2-Back – participants pressed whenever the letter presented matched the one shown two letters prior.

*WASI-II Matrix Reasoning.* In this abstract reasoning task, participants were presented with an incomplete matrix and asked to select from a choice of four response options the item that best completed each matrix.

*WASI-II Block Design.* In this reasoning task, participants were instructed to use red and white blocks to reconstruct designs of increasing difficulty.

*WASI-II Similarities.* In this verbal reasoning task, participants were asked to identify items sharing similarities with other items or to describe in what way(s) certain words were similar.

*CNB Verbal Reasoning.* Participants completed multiple-choice verbal analogy problems.

*DKEFS Trail-Making.* This motor-speed task, which taps into both clerical speed and choice reaction time components, participants are asked to draw a line to link circles as quickly and accurately as possible according to specified rules (e.g. numbers in ascending order such as 1-2-3…, or switching between numbers and letters such as 1-A-2-B…). Participants' scores reflected the total completion time.

*PhAB Rapid Naming.* In this clerical speed task, participants were shown a card displaying commonplace objects (hat, table, box, ball, and door) and asked to name them as quickly and accurately as possible. The total time taken to complete the task was modeled.

*TEA-Ch RBBS.* In this choice reaction time and switching task, participants were presented with items (e.g. a red handbag) and instructed to either match them to the right colour (red or blue) or use (hand or foot). Participants' scores were calculated from the average response time for each trial.

*CNB Motor Speed.* Participants were asked to tap a key as quickly and as many times as possible within 10 seconds in this clerical speed task. The total number of taps was modeled.

*CNB Sensory Motor Speed.* Participants were shown a moving green square on a screen and asked to click on it as rapidly as possible in this clerical speed task. Median response time was used as the performance score.

### White Matter Microstructure.

*MRI data acquisition.* Magnetic resonance imaging data were acquired at the MRC Cognition and Brain Sciences Unit, Cambridge U.K. for CALM. All scans for CALM were obtained on the Siemens 3 T Tim Trio system (Siemens Healthcare, Erlangen, Germany), using a 32-channel quadrature head coil. Diffusion scans were acquired using echo-planar diffusion-weighted images with a set of 60 non-collinear directions, using a weighting factor of b=1000s*mm$^{-2}$, interleaved with a T2-weighted (b=0) volume. Whole brain coverage was obtained with 60 contiguous axial slices and isometric image resolution of 2mm. Echo time was 90ms and repetition time was 8400ms. NKI-RS participants were assessed using a Siemens TrioTM 3 T MRI scanner. T1-weighted images were acquired using a magnetization-prepared rapid gradient echo (MPRAGE) sequence with 1mm isotropic resolution. Diffusion scans were acquired with 137-direction, 2mm isotropic sequence using a weighting factor of b=1000s*mm$^{-2}$. Details of the NKI-RS scan sequences can be found here: http://fcon_1000.projects.nitrc.org/indi/enhanced/mri_protocol.html.

*MRI quality control.* Participant movement may significantly affect the quality of MRI data and may bias statistical comparisons, especially in developmental populations (Power et al. 2012; Savalia et al. 2016). Several steps were taken to assure good MRI data quality and minimize potential biases of participant movement. First, children were instructed to lie still and were trained to do so in a realistic mock scanner prior to the actual scan for CALM. Second, all T1-weighted images and FA maps for both samples were visually inspected by a trained researcher (J.B.) to remove low quality scans. Further, the quality of the diffusion-weighted data were assessed by calculating the displacement between subsequent volumes in the sequence (see below). Only DWI data with between-volume displacement below 3mm were included in the analysis. For the included datasets, the maximum between-volume displacement was used a covariate to control for spurious effects associated with participant movement (see Testing for potential confounds).

*Processing of DTI data.* Diffusion-weighted images were pre-processed to create a brain mask based on the b0-weighted image (FSL BET) (Smith 2002) and to correct for movement and eddy current-induced distortions (*eddy*) (Graham et al. 2016). Subsequently, the diffusion tensor model was fitted and FA maps were calculated (*dtifit*). Images with a between-image displacement great than 3mm as indicated by FSL eddy were excluded from further analysis. All steps were carried out with FSL v5.0.9 and were implemented in a pipeline using NiPyPe v0.13.0 (Gorgolewski et al. 2011). To extract FA values for major white matter tracts, FA images were registered to the FMRIB58 FA template in MNI space using a sequence of rigid co-registration, followed by affine co-registration, and symmetric diffeomorphic image registration (*SyN*) as implemented in ANTS v1.9 (Avants et al. 2008). Visual inspection indicated good image registration for all participants. Subsequently, binary masks from the JHU white matter atlas in MNI space (Mori et al. 2008) were applied to extract FA values for 10 major white matter tracts bilaterally.

### Supplementary Analyses

**Testing for Potential Confounds**. We carried out a series of supplementary analyses to examine whether possible confounders influenced our models. These analyses showed that our inferences were robust to the inclusion of covariates such as scanner motion or socio-economic status. They were also robust across genders, and participants taking or not taking medication, while differences cannot be ruled out for participants with and without diagnosed disorders in NKI-RS.

To test confounding by continuous variables, we included these variables as covariates in our models. To test confounding by categorical variables, we ran multi-group models and imposed a set of canonical, increasingly restrictive invariance constraints (Widaman and Reise 1997; Steenkamp and Baumgartner 1998). First we established configural invariance by fitting a freely-estimated model and examining model fit. This tests the hypothesis that the same overall model fits for both groups. Next, we tested weak invariance but constraining factor loadings to be equal across groups. This weak invariance model was then compared to the configural invariance model to test the hypothesis that the loadings differ significantly between groups. Finally, we tested strong invariance by constraining factor loadings and intercepts to be equal across groups. This model was then compared to the weak-invariance model to test the hypothesis that loadings and intercepts differ significantly between groups. Overall, these equality constraints test the hypothesis that although there are small differences between the cohorts, the overall watershed model holds.

*Scanner Motion.* To test whether motion in the scanner confounded results, we re-ran models including a covariate for maximum frame-wise displacement. Specifically, the watershed model included additional regression paths from the motion parameter to FA in all tracts. Model fit was good for CALM ($\chi^2(86) = 134.36$, $p < .001$; RMSEA = .032 [.021 - .042]; CFI = .984; SRMR = .052) and acceptable fit for NKI-RS ($\chi^2(122) = 239.56$, $p < .001$; RMSEA = .054 [.043 - .064]; CFI = .953; SRMR = .103) and all originally significant paths retained their directionality and significance (Supplementary Table 5; Supplementary Table 6).

*Socio-economic status.* To assess the impact of socio-economic status we modeled household income level standardized by the number of people living in the household for NKI-RS. For CALM, there is currently no data on socio-economic status available. Socio-economic status was included as an additional predictor of memory, speed and $g_f$. We found that the regression paths in the NKI-RS model changed very little (Supplementary Table 7) and model fit remained very similar ($\chi^2(119) = 231.91$, $p < .001$; RMSEA = .053 [.042 - .064]; CFI = .926; SRMR = .092).

*Gender differences.* Multi-group models of the NKI-RS watershed model showed convergence problems, likely due to splitting the already relatively small number of participants with neuroimaging data ($N = 67$) into subgroups. We therefore analysed the MIMIC model without neuroimaging data for this sample. For gender, a model imposing configural invariance constraints fit well for CALM ($\chi^2(156) = 202.15$, $p = .008$; RMSEA = .033 [.017 - .045]; CFI = .971; SRMR = .053) while it did not converge for NKI-RS, likely still due to the low sample size in each group ($N_{\text{female}} = 149$). The weak invariance model showed acceptable fit for NKI-RS ($\chi^2(64) = 72.93$, $p = .208$; RMSEA = .055 [.000 - .107]; CFI = .974; SRMR = .060). For CALM we found no statistical differences between a model with configural and weak ($\Delta\chi^2(8) = 4.30$, $p = .829$) and weak and strong invariance constraints across sexes ($\Delta\chi^2(5) = 0.19$, $p = .999$), suggesting that a model with strong invariance constraints was most parsimonious. There was also no difference between weak and strong invariance constraints for NKI-RS ($\Delta\chi^2(7) = 2.15$, $p = .951$). This indicates that the overall model structure, factor loadings and intercepts did not differ significantly between males and females for both samples.

*Diagnosed disorders.* We tested whether our models differed across participants with and without confirmed diagnoses (See Table 1) following the same procedure of invariance tests described above. For CALM, the configural invariance model fit well ($\chi^2(156) = 210.80$, $p = .002$; RMSEA = .036 [.022 - .048]; CFI = .965; SRMR = .062). We found no statistical differences between a model with configural and weak ($\Delta\chi^2(8) = 5.27$, $p = .729$) and weak and strong invariance constraints ($\Delta\chi^2(5) = 1.49$, $p = .915$), suggesting that a model with strong invariance constraints was most parsimonious. For NKI-RS the configural invariance model again failed to converge. The weak invariance model showed poor fit in NKI-RS ($\chi^2(234) = 1066.02$, $p < .001$; RMSEA = .146 [.135 - .157]; CFI = .670; SRMR = .136). This is likely be due to the low number of participants with diagnosed disorders ($N = 106$), but means that we cannot rule out potential group differences within NKI-RS (Supplementary Table 8).

*Medication.* We tested whether our models differed across participants taking or not taking medication in NKI-RS and participants taking or not talking ADHD medication in CALM. For

CALM and NKI-RS, the full watershed model did not converge, likely due to the low number of medicated participants ($N_{CALM}$ = 58, $N_{NKI\text{-}RS}$ = 57). We therefore tested invariance in the MIMC model for both samples. For CALM, the configural invariance model fit well ($\chi^2$(36) = 61.52, $p$ = .005; RMSEA = .052 [.027 - .074]; CFI = .982; SRMR = .036, and we found no statistical differences between a model with configural and weak ($\Delta\chi^2$(8) = 6.01, $p$ = .647) and weak and strong invariance constraints ($\Delta\chi^2$(5) = 0.58, $p$ = .989), suggesting that a model with strong invariance constraints was most parsimonious. For NKI-R, the configural invariance model yielded an improper solution but the weak invariance model showed acceptable fit ($\chi^2$(74) = 120.87, p < .001; RMSEA = .064 [.042 - .084]; CFI = .960; SRMR = .052). There was no difference between a model with weak and a strong invariance constrains ($\Delta\chi^2$ (7) = 0.94, $p$ = .996), suggesting that a model with strong invariance constraints was most parsimonious for NKI-RS. This indicates that the overall MIMC model structure, factor loadings and intercepts did not differ significantly between participants taking or not taking medications for both samples.

### Supplementary Figures

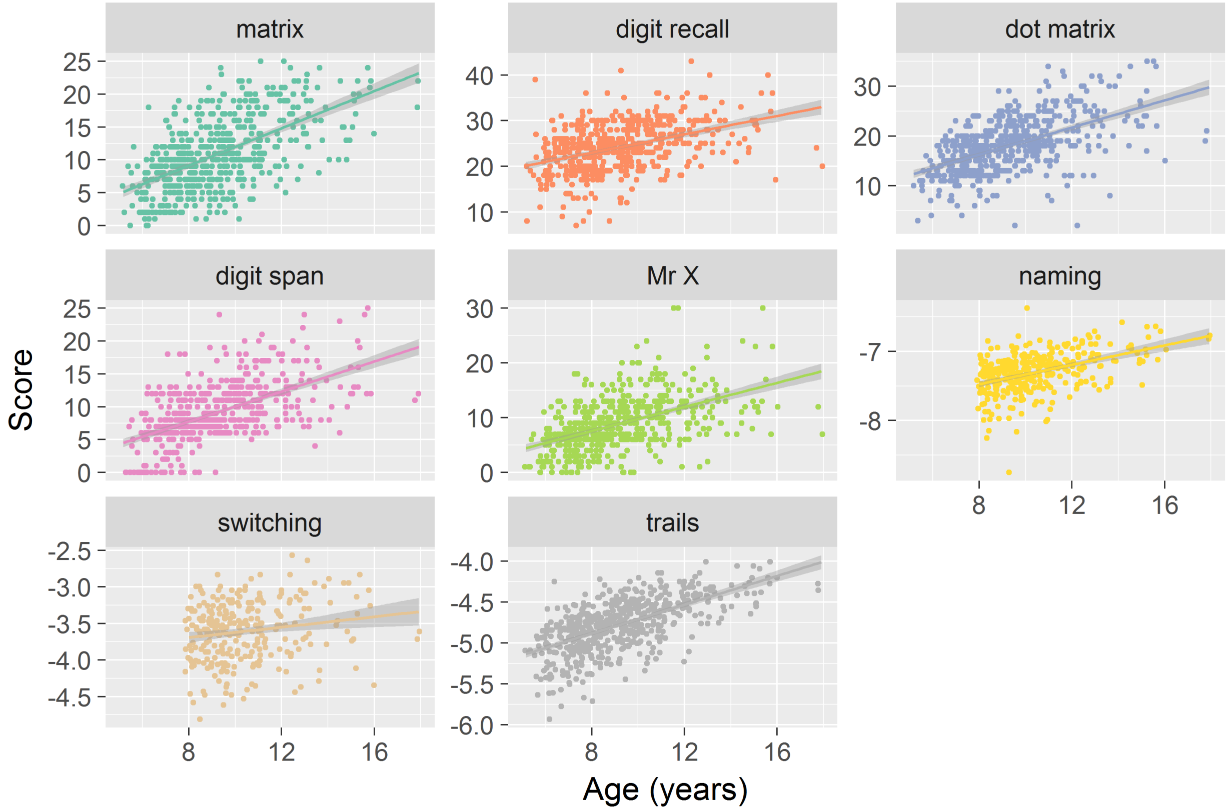


*Supplementary Figure 1. Scatterplots of Task Scores against Age for CALM.*

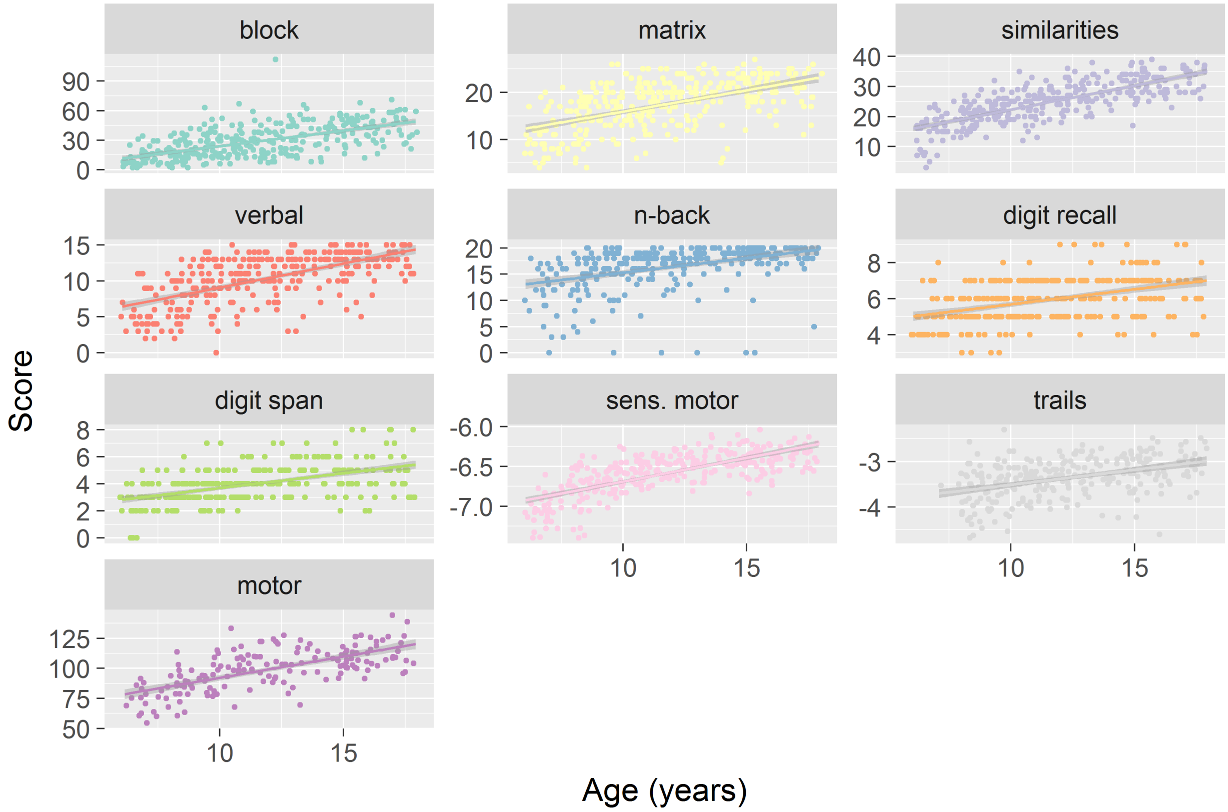


168

169 *Supplementary Figure 2. Scatterplots of Task Scores against Age for NKI.*

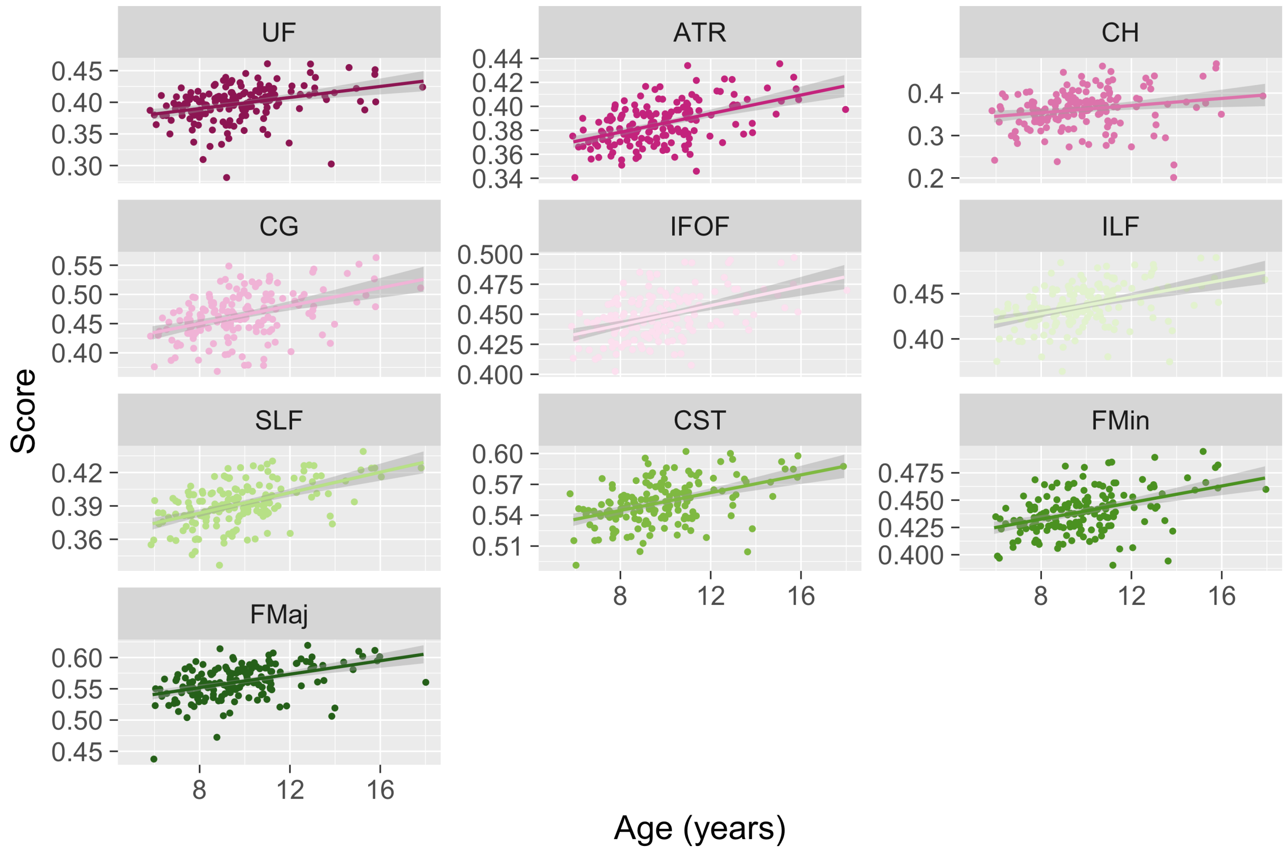


*Supplementary Figure 3. Scatterplots of FA in the 10 White Matter Tracts against Age for CALM.* Abbreviations: uncinate fasciculus (UF), superior longitudinal fasciculus (SLF), inferior fronto-occipital fasciculus (IFOF), anterior thalamic radiations (ATR), cerebrospinal tract (CST), forceps major (FMaj), forceps minor (FMin), dorsal cingulate gyrus (CG), ventral cingulate gyrus (CH), inferior longitudinal fasciculus (ILF).

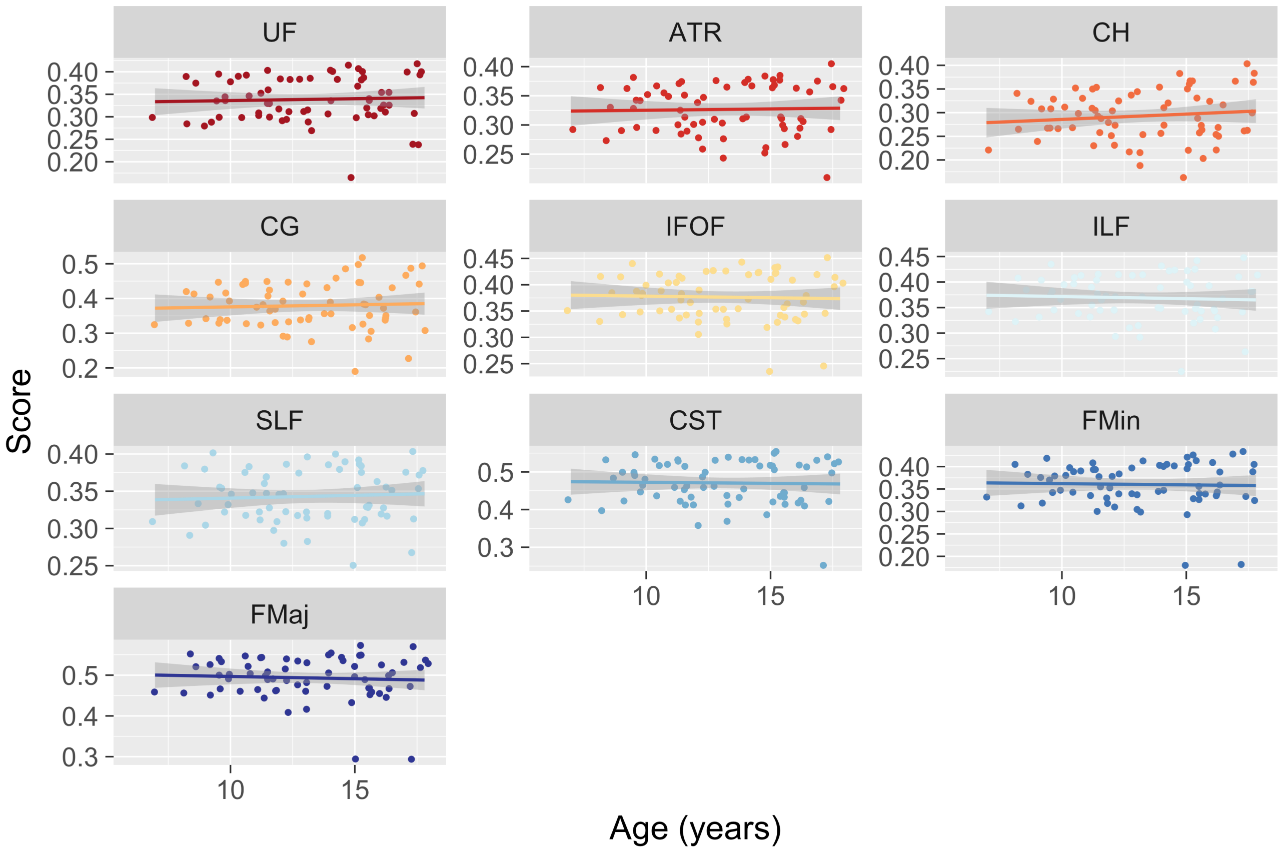


*Supplementary Figure 4. Scatterplots of FA in the 10 White Matter Tracts against Age for NKI.*

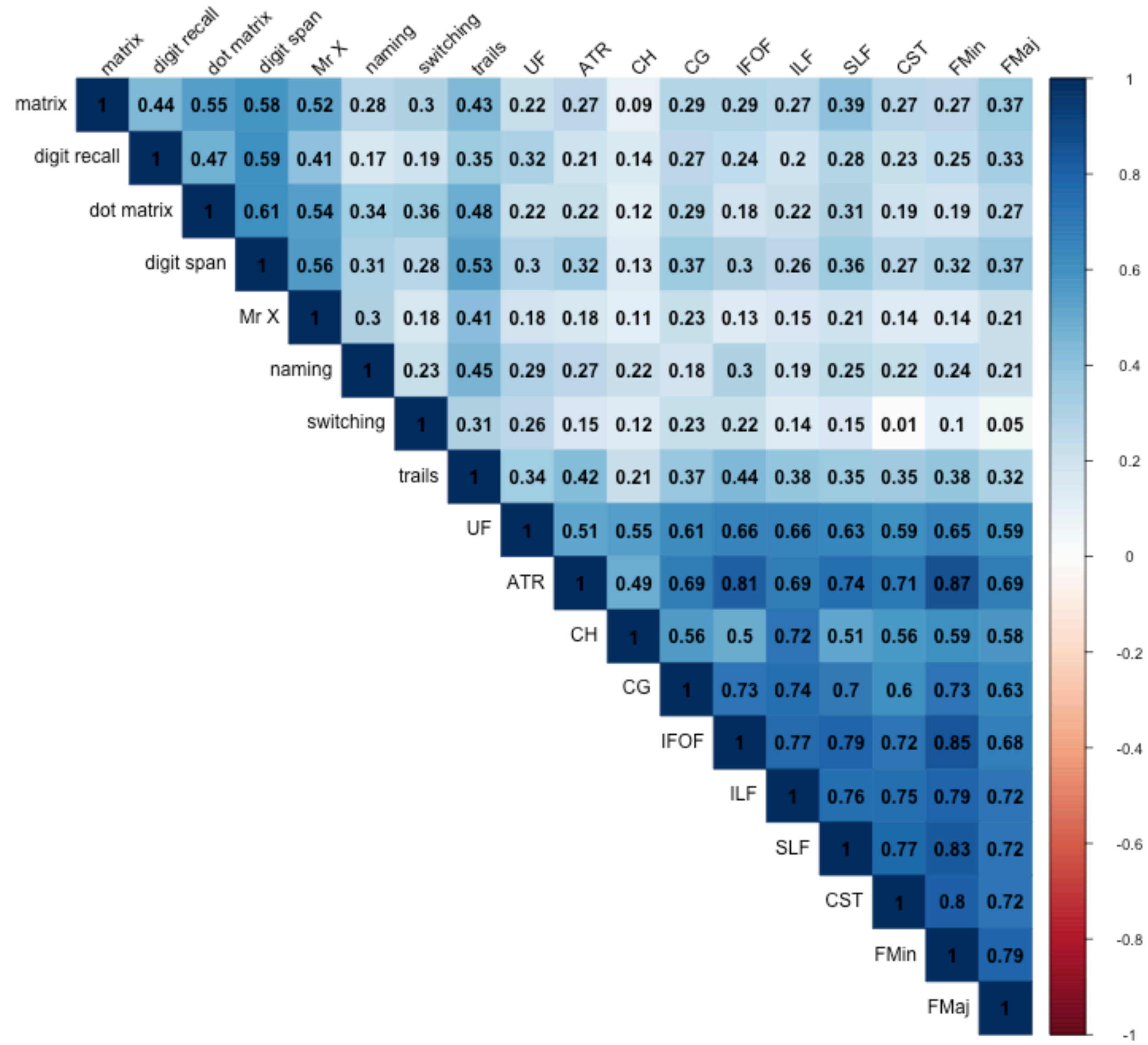


*Supplementary Figure 5. Correlation Matrix of Tasks and White Matter Tracts Modeled in CALM.*

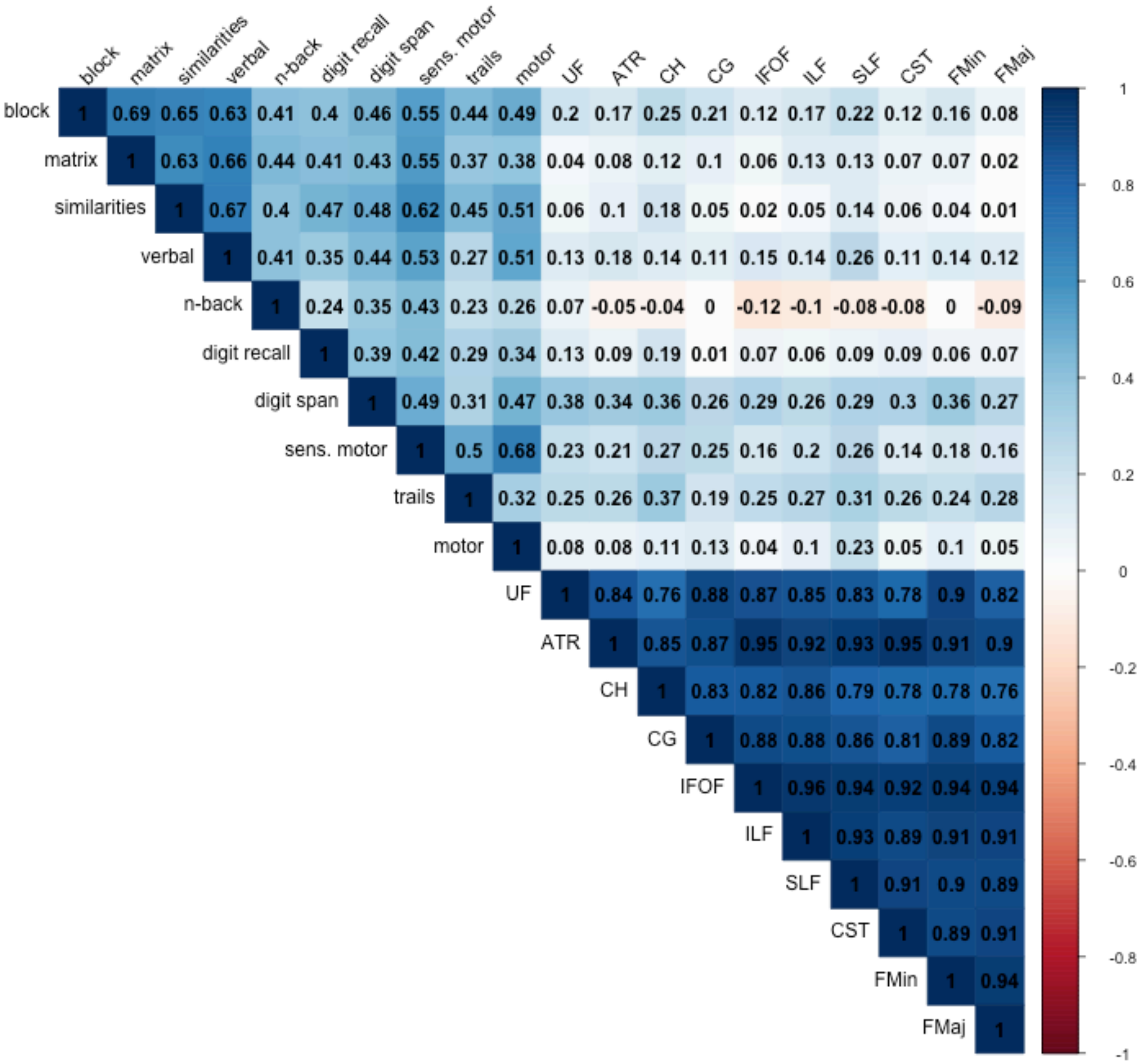


*Supplementary Figure 6. Correlation Matrix of Tasks and White Matter Tracts Modeled in NKI.*

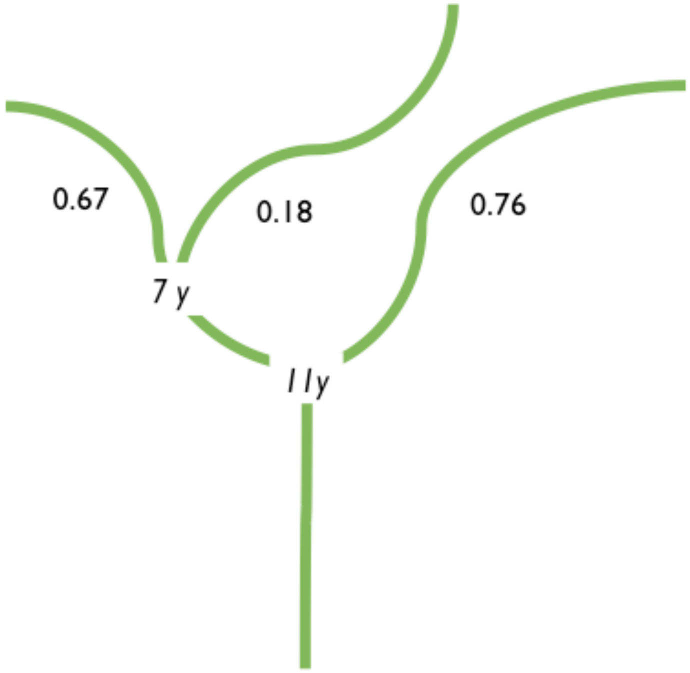


*Supplementary Figure 7. Example for an SEM Tree: The Relationship between the SLF and memory in CALM.* The two age splits in years are shown in cursive, with parameter estimates before and after each split displayed above. See Table 6 for all other SEM tree results.

### Supplementary Tables

*Supplementary Table 1. Factor Loadings of the Three-Factor Measurement Model*

| Path | | | Estimate | SE | z | p | Standardized Estimate |
|---|---|---|---|---|---|---|---|
| | | | CALM | | | | |
| $g_f$ | =~ | matrix | 1.00 | 0.03 | 38.80 | < .001 | 1.00 |
| memory | =~ | digit recall | 0.64 | 0.04 | 14.74 | < .001 | 0.64 |
| memory | =~ | dot-matrix | 0.76 | 0.04 | 18.42 | < .001 | 0.76 |
| memory | =~ | digit span | 0.83 | 0.04 | 19.36 | < .001 | 0.83 |
| memory | =~ | Mr X | 0.69 | 0.04 | 15.41 | < .001 | 0.69 |
| speed | =~ | PhAB | 0.79 | 0.05 | 15.82 | < .001 | 0.79 |
| speed | =~ | Teach | 0.66 | 0.07 | 9.05 | < .001 | 0.64 |
| speed | =~ | Trails | 0.53 | 0.08 | 7.09 | < .001 | 0.51 |
| | | | NKI-RS | | | | |
| $g_f$ | =~ | block | 0.81 | 0.04 | 22.25 | < .001 | 0.81 |
| $g_f$ | =~ | matrix | 0.81 | 0.04 | 20.05 | < .001 | 0.81 |
| $g_f$ | =~ | similarities | 0.82 | 0.05 | 18.12 | < .001 | 0.82 |
| $g_f$ | =~ | verbal | 0.81 | 0.04 | 19.23 | < .001 | 0.81 |
| memory | =~ | n-back | 0.58 | 0.05 | 10.58 | < .001 | 0.57 |
| memory | =~ | digit recall | 0.54 | 0.06 | 9.12 | < .001 | 0.55 |
| memory | =~ | digit span | 0.62 | 0.07 | 9.42 | < .001 | 0.62 |
| speed | =~ | sens. motor | 0.88 | 0.05 | 17.43 | < .001 | 0.88 |
| speed | =~ | Trails | 0.74 | 0.07 | 10.22 | < .001 | 0.69 |
| speed | =~ | motor | 0.81 | 0.07 | 11.72 | < .001 | 0.78 |

*Supplementary Table 2. Regression Paths of the Watershed Model for CALM*

| Path | | | Estimate | SE | *z* | *p* | Standardized Estimate |
|---|---|---|---|---|---|---|---|
| $g_f$ | ~ | speed | -0.03 | 0.11 | -0.29 | .768 | -0.03 |
| $g_f$ | ~ | memory | 0.87 | 0.12 | 7.14 | < .001 | 0.74 |
| speed | ~ | UF | 0.28 | 0.16 | 1.82 | .069 | 0.22 |
| speed | ~ | SLF | -0.02 | 0.20 | -0.10 | .919 | -0.02 |
| speed | ~ | IFOF | 0.37 | 0.23 | 1.61 | .108 | 0.29 |
| speed | ~ | ATR | 0.49 | 0.23 | 2.08 | .038 | 0.39 |
| speed | ~ | CST | 0.06 | 0.20 | 0.30 | .768 | 0.05 |
| speed | ~ | FMaj | -0.01 | 0.19 | -0.04 | .966 | -0.01 |
| speed | ~ | FMin | -0.47 | 0.33 | -1.43 | .152 | -0.37 |
| speed | ~ | CG | 0.11 | 0.14 | 0.78 | .435 | 0.09 |
| speed | ~ | CH | -0.11 | 0.17 | -0.62 | .534 | -0.08 |
| speed | ~ | ILF | 0.14 | 0.23 | 0.59 | .553 | 0.11 |
| memory | ~ | UF | 0.21 | 0.13 | 1.58 | .114 | 0.17 |
| memory | ~ | SLF | 0.43 | 0.17 | 2.54 | .011 | 0.35 |
| memory | ~ | IFOF | -0.15 | 0.20 | -0.77 | .440 | -0.13 |
| memory | ~ | ATR | 0.23 | 0.19 | 1.23 | .218 | 0.19 |
| memory | ~ | CST | -0.05 | 0.16 | -0.32 | .746 | -0.04 |
| memory | ~ | FMaj | 0.47 | 0.13 | 3.62 | < .001 | 0.39 |
| memory | ~ | FMin | -0.51 | 0.25 | -2.02 | .043 | -0.42 |
| memory | ~ | CG | 0.34 | 0.12 | 2.72 | .007 | 0.28 |
| memory | ~ | CH | -0.22 | 0.16 | -1.37 | .170 | -0.18 |
| memory | ~ | ILF | -0.09 | 0.19 | -0.47 | .639 | -0.07 |

*Supplementary Table 3. Regression Paths of the Watershed Model for NKI-RS*

| Path | | | Estimate | SE | *z* | *p* | Standardized Estimate |
|---|---|---|---|---|---|---|---|
| $g_f$ | ~ | speed | 0.42 | 0.35 | 1.22 | .224 | 0.29 |
| $g_f$ | ~ | memory | 0.99 | 0.63 | 1.56 | .119 | 0.63 |
| speed | ~ | UF | 0.03 | 0.62 | 0.04 | .966 | 0.02 |
| speed | ~ | SLF | 2.37 | 0.75 | 3.17 | .002 | 1.67 |
| speed | ~ | IFOF | -2.45 | 0.90 | -2.73 | .006 | -1.67 |
| speed | ~ | ATR | 0.13 | 0.75 | 0.18 | .858 | 0.09 |
| speed | ~ | CST | -0.83 | 0.61 | -1.35 | .177 | -0.56 |
| speed | ~ | FMaj | 0.38 | 0.75 | 0.51 | .610 | 0.26 |
| speed | ~ | FMin | 0.35 | 0.78 | 0.45 | .652 | 0.24 |
| speed | ~ | CG | 0.01 | 0.53 | 0.02 | .984 | 0.01 |
| speed | ~ | CH | 0.76 | 0.44 | 1.71 | .087 | 0.53 |
| speed | ~ | ILF | -0.18 | 0.74 | -0.25 | .806 | -0.13 |
| memory | ~ | UF | 0.53 | 0.56 | 0.94 | .349 | 0.39 |
| memory | ~ | SLF | 1.82 | 0.59 | 3.10 | .002 | 1.39 |
| memory | ~ | IFOF | -2.14 | 0.92 | -2.33 | .020 | -1.58 |
| memory | ~ | ATR | 0.35 | 0.78 | 0.45 | .654 | 0.26 |
| memory | ~ | CST | -0.32 | 0.62 | -0.52 | .603 | -0.24 |
| memory | ~ | FMaj | -0.76 | 0.59 | -1.27 | .203 | -0.56 |
| memory | ~ | FMin | 1.16 | 0.70 | 1.65 | .099 | 0.86 |
| memory | ~ | CG | -0.68 | 0.54 | -1.25 | .210 | -0.51 |
| memory | ~ | CH | 0.82 | 0.43 | 1.92 | .055 | 0.63 |
| memory | ~ | ILF | -0.48 | 0.69 | -0.69 | .493 | -0.36 |

*Supplementary Table 4. SEM Tree Splits with a Maximum Depth of One Split*

| Path | Estimate Before | Age Split | Estimate After |
|---|---|---|---|
| | *CALM* | | |
| memory <–> speed | 0.89 | 9.46 | 0.74 |
| memory –> $g_f$ | 0.80 | 10.04 | 0.97 |
| speed –> $g_f$ | -0.11 | 11.21 | 0.17 |
| SLF –> memory | 0.31 | 11.21 | 0.76 |
| FMaj –> memory | 0.39 | 9.29 | 0.59 |
| CG –> memory | 0.20 | 11.04 | 0.70 |
| ATR –> speed | 0.85 | 7.96 | 0.33 |
| | *NKI-RS* | | |
| memory <–> speed | 0.73 | 14.72 | 1.11 |
| memory –> $g_f$ | 1.10 | 8.59 | 0.87 |
| speed –> $g_f$ | 0.53 | 8.59 | 0.27 |
| SLF –> memory | 2.15 | 8.30 | 1.70 |
| SLF –> speed | 3.12 | 8.63 | 2.00 |

*Supplementary Table 5. Regression Paths of the Watershed Model for CALM Including Scanner Motion (FWD) as a Covariate*

| Path | | | Estimate | SE | *z* | *p* | Standardized Estimate |
|---|---|---|---|---|---|---|---|
| $g_f$ | ~ | speed | -0.03 | 0.11 | -0.29 | .768 | -0.03 |
| $g_f$ | ~ | memory | 0.87 | 0.12 | 7.14 | < .001 | 0.74 |
| speed | ~ | UF | 0.28 | 0.16 | 1.82 | .069 | 0.22 |
| speed | ~ | SLF | -0.02 | 0.20 | -0.10 | .919 | -0.02 |
| speed | ~ | IFOF | 0.37 | 0.23 | 1.61 | .108 | 0.29 |
| speed | ~ | ATR | 0.49 | 0.23 | 2.08 | .038 | 0.39 |
| speed | ~ | CST | 0.06 | 0.20 | 0.30 | .768 | 0.05 |
| speed | ~ | FMaj | -0.01 | 0.19 | -0.04 | .966 | -0.01 |
| speed | ~ | FMin | -0.47 | 0.33 | -1.43 | .152 | -0.37 |
| speed | ~ | CG | 0.11 | 0.14 | 0.78 | .435 | 0.09 |
| speed | ~ | CH | -0.11 | 0.17 | -0.62 | .534 | -0.08 |
| speed | ~ | ILF | 0.14 | 0.23 | 0.59 | .553 | 0.11 |
| memory | ~ | UF | 0.21 | 0.13 | 1.58 | .114 | 0.17 |
| memory | ~ | SLF | 0.43 | 0.17 | 2.54 | .011 | 0.35 |
| memory | ~ | IFOF | -0.15 | 0.20 | -0.77 | .440 | -0.13 |
| memory | ~ | ATR | 0.23 | 0.19 | 1.23 | .218 | 0.19 |
| memory | ~ | CST | -0.05 | 0.16 | -0.32 | .746 | -0.04 |
| memory | ~ | FMaj | 0.47 | 0.13 | 3.62 | < .001 | 0.39 |
| memory | ~ | FMin | -0.51 | 0.25 | -2.02 | .043 | -0.42 |
| memory | ~ | CG | 0.34 | 0.12 | 2.72 | .007 | 0.28 |
| memory | ~ | CH | -0.22 | 0.16 | -1.37 | .170 | -0.18 |
| memory | ~ | ILF | -0.09 | 0.19 | -0.47 | .639 | -0.07 |
| UF | ~ | motion | -0.08 | 0.08 | -1.10 | .273 | -0.08 |
| SLF | ~ | motion | -0.09 | 0.07 | -1.27 | .205 | -0.09 |
| IFOF | ~ | motion | -0.04 | 0.07 | -0.52 | .601 | -0.04 |
| ATR | ~ | motion | 0.01 | 0.08 | 0.16 | .875 | 0.01 |
| CST | ~ | motion | -0.07 | 0.07 | -0.91 | .364 | -0.06 |
| FMaj | ~ | motion | -0.09 | 0.07 | -1.22 | .221 | -0.09 |
| FMin | ~ | motion | 0.02 | 0.07 | 0.32 | .746 | 0.02 |
| CG | ~ | motion | -0.05 | 0.08 | -0.62 | .534 | -0.05 |
| CH | ~ | motion | -0.12 | 0.07 | -1.64 | .102 | -0.12 |
| ILF | ~ | motion | -0.15 | 0.07 | -2.04 | .041 | -0.15 |

*Supplementary Table 6. Regression Paths of the Watershed Model for NKI-RS Including Scanner Motion (FWD) as a Covariate*

| Path | | | Estimate | SE | *z* | *p* | Standardized Estimate |
|---|---|---|---|---|---|---|---|
| $g_f$ | ~ | speed | 0.42 | 0.34 | 1.22 | .224 | 0.29 |
| $g_f$ | ~ | memory | 0.99 | 0.63 | 1.56 | .119 | 0.63 |
| speed | ~ | UF | 0.03 | 0.62 | 0.04 | .966 | 0.02 |
| speed | ~ | SLF | 2.37 | 0.75 | 3.17 | .002 | 1.67 |
| speed | ~ | IFOF | -2.45 | 0.90 | -2.73 | .006 | -1.67 |
| speed | ~ | ATR | 0.13 | 0.75 | 0.18 | .858 | 0.09 |
| speed | ~ | CST | -0.83 | 0.61 | -1.35 | .177 | -0.56 |
| speed | ~ | FMaj | 0.38 | 0.75 | 0.51 | .610 | 0.26 |
| speed | ~ | FMin | 0.35 | 0.78 | 0.45 | .652 | 0.24 |
| speed | ~ | CG | 0.01 | 0.53 | 0.02 | .984 | 0.01 |
| speed | ~ | CH | 0.76 | 0.44 | 1.71 | .087 | 0.53 |
| speed | ~ | ILF | -0.18 | 0.74 | -0.25 | .806 | -0.13 |
| memory | ~ | UF | 0.53 | 0.56 | 0.94 | .349 | 0.39 |
| memory | ~ | SLF | 1.82 | 0.59 | 3.10 | .002 | 1.39 |
| memory | ~ | IFOF | -2.14 | 0.92 | -2.33 | .020 | 1.58 |
| memory | ~ | ATR | 0.35 | 0.78 | 0.45 | .654 | 0.26 |
| memory | ~ | CST | -0.32 | 0.62 | -0.52 | .604 | -0.24 |
| memory | ~ | FMaj | -0.76 | 0.59 | -1.27 | .203 | -0.56 |
| memory | ~ | FMin | 1.16 | 0.70 | 1.65 | .099 | 0.86 |
| memory | ~ | CG | -0.68 | 0.54 | -1.25 | .209 | -0.51 |
| memory | ~ | CH | 0.82 | 0.43 | 1.92 | .055 | 0.63 |
| memory | ~ | ILF | -0.48 | 0.69 | -0.69 | .493 | -0.36 |
| UF | ~ | motion | -0.09 | 0.12 | -0.71 | .476 | -0.09 |
| SLF | ~ | motion | -0.15 | 0.11 | -1.38 | .169 | -0.15 |
| IFOF | ~ | motion | -0.11 | 0.15 | -0.73 | .463 | -0.11 |
| ATR | ~ | motion | -0.14 | 0.14 | -0.99 | .324 | -0.13 |
| CST | ~ | motion | -0.16 | 0.19 | -0.85 | .393 | -0.16 |
| FMaj | ~ | motion | -0.20 | 0.18 | -1.10 | .271 | -0.20 |
| FMin | ~ | motion | -0.16 | 0.18 | -0.92 | .360 | -0.16 |
| CG | ~ | motion | -0.16 | 0.13 | -1.23 | .220 | -0.15 |
| CH | ~ | motion | 0.01 | 0.10 | 0.09 | .926 | 0.01 |
| ILF | ~ | motion | -0.05 | 0.14 | -0.39 | .695 | -0.05 |

*Supplementary Table 7. Regression Paths of the Watershed Model for NKI-RS Including SES as a Covariate*

| Path | | | Estimate | SE | *z* | *p* | Standardized Estimate |
|---|---|---|---|---|---|---|---|
| $g_f$ | ~ | speed | 0.42 | 0.33 | 1.26 | .207 | 0.30 |
| $g_f$ | ~ | memory | 0.91 | 0.61 | 1.51 | .131 | 0.61 |
| $g_f$ | ~ | SES | 0.13 | 0.09 | 1.51 | .132 | 0.06 |
| speed | ~ | UF | -0.10 | 0.63 | -0.16 | .869 | -0.07 |
| speed | ~ | SLF | 2.41 | 0.82 | 2.94 | .003 | 1.64 |
| speed | ~ | IFOF | -2.30 | 0.87 | -2.64 | .008 | -1.52 |
| speed | ~ | ATR | 0.42 | 0.87 | 0.49 | .626 | 0.28 |
| speed | ~ | CST | -1.56 | 0.91 | -1.71 | .087 | -1.02 |
| speed | ~ | FMaj | 0.56 | 0.75 | 0.75 | .456 | 0.37 |
| speed | ~ | FMin | 0.78 | 0.83 | 0.94 | .349 | 0.52 |
| speed | ~ | CG | -0.12 | 0.58 | -0.21 | .835 | -0.08 |
| speed | ~ | CH | 0.73 | 0.43 | 1.70 | .089 | 0.49 |
| speed | ~ | ILF | -0.24 | 0.78 | -0.31 | .760 | -0.16 |
| speed | ~ | SES | 0.32 | 0.26 | 1.25 | .212 | 0.21 |
| memory | ~ | UF | 0.43 | 0.58 | 0.74 | .456 | 0.31 |
| memory | ~ | SLF | 1.84 | 0.66 | 2.79 | .005 | 1.34 |
| memory | ~ | IFOF | -1.93 | 0.88 | -2.19 | .029 | -1.37 |
| memory | ~ | ATR | 0.66 | 0.92 | 0.71 | .476 | 0.47 |
| memory | ~ | CST | -1.16 | 0.87 | -1.33 | .182 | -0.82 |
| memory | ~ | FMaj | -0.59 | 0.60 | -0.97 | .331 | -0.42 |
| memory | ~ | FMin | 1.72 | 0.82 | 2.08 | .037 | 1.23 |
| memory | ~ | CG | -0.88 | 0.58 | -1.51 | .132 | -0.63 |
| memory | ~ | CH | 0.81 | 0.42 | 1.91 | .056 | 0.59 |
| memory | ~ | ILF | -0.58 | 0.77 | -0.75 | .451 | -0.42 |
| memory | ~ | SES | 0.38 | 0.24 | 1.60 | .110 | 0.27 |

*Supplementary Table 8. Regression Paths of the Watershed Model for NKI-RS for Participants with and without a Diagnosed Disorder.*

| Path | | | Estimate | SE | *z* | *p* | Standardized Estimate |
|---|---|---|---|---|---|---|---|
| | | | *Without diagnosis (N = 229)* | | | | |
| $g_f$ | ~ | speed | 0.42 | 0.24 | 1.76 | .079 | 0.36 |
| $g_f$ | ~ | memory | 0.70 | 0.46 | 1.52 | .129 | 0.55 |
| speed | ~ | UF | 0.45 | 0.34 | 1.31 | .192 | 0.26 |
| speed | ~ | SLF | 3.59 | 0.73 | 4.93 | <.001 | 2.09 |
| speed | ~ | IFOF | -3.16 | 1.01 | -3.13 | .002 | -1.80 |
| speed | ~ | ATR | 0.32 | 1.18 | 0.27 | .786 | 0.19 |
| speed | ~ | CST | -0.55 | 0.83 | -0.66 | .513 | -0.31 |
| speed | ~ | FMaj | 0.40 | 0.71 | 0.57 | .566 | 0.23 |
| speed | ~ | FMin | 0.85 | 0.75 | 1.13 | .259 | 0.48 |
| speed | ~ | CG | -0.47 | 0.76 | -0.62 | .535 | -0.27 |
| speed | ~ | CH | 1.28 | 0.51 | 2.49 | .013 | 0.75 |
| speed | ~ | ILF | -1.87 | 0.88 | -2.13 | .033 | -1.07 |
| memory | ~ | UF | 1.54 | 1.07 | 1.43 | .152 | 0.97 |
| memory | ~ | SLF | 1.76 | 1.06 | 1.67 | .095 | 1.12 |
| memory | ~ | IFOF | -2.33 | 1.62 | -1.44 | .150 | -1.45 |
| memory | ~ | ATR | 0.95 | 1.68 | 0.57 | .572 | 0.60 |
| memory | ~ | CST | -0.79 | 1.05 | -0.75 | .453 | -0.49 |
| memory | ~ | FMaj | -1.03 | 1.01 | -1.02 | .308 | -0.64 |
| memory | ~ | FMin | 0.83 | 1.25 | 0.67 | .506 | 0.52 |
| memory | ~ | CG | -1.25 | 0.94 | -1.33 | .185 | -0.78 |
| memory | ~ | CH | 0.74 | 0.74 | 1.00 | .319 | 0.47 |
| memory | ~ | ILF | -0.13 | 1.23 | -0.10 | .917 | -0.08 |
| | | | *With diagnosis (N = 106)* | | | | |
| $g_f$ | ~ | speed | 1.06 | 0.31 | 3.46 | .001 | 0.86 |
| $g_f$ | ~ | memory | 0.01 | 0.19 | 0.07 | .945 | 0.01 |
| speed | ~ | UF | -0.85 | 1.11 | -0.77 | .444 | -0.53 |
| speed | ~ | SLF | 1.04 | 0.72 | 1.44 | .149 | 0.70 |
| speed | ~ | IFOF | -1.29 | 1.68 | -0.77 | .444 | -0.81 |
| speed | ~ | ATR | 0.57 | 1.44 | 0.40 | .690 | 0.37 |
| speed | ~ | CST | -0.70 | 0.82 | -0.85 | .394 | -0.45 |
| speed | ~ | FMaj | 0.05 | 1.27 | 0.04 | .968 | 0.03 |
| speed | ~ | FMin | 0.77 | 1.65 | 0.47 | .639 | 0.51 |
| speed | ~ | CG | 1.00 | 0.84 | 1.18 | .237 | 0.68 |
| speed | ~ | CH | -0.28 | 0.72 | -0.39 | .699 | -0.17 |
| speed | ~ | ILF | 0.46 | 0.91 | 0.51 | .612 | 0.30 |
| memory | ~ | UF | -0.56 | 1.21 | -0.46 | .646 | -0.34 |
| memory | ~ | SLF | 1.43 | 0.93 | 1.54 | .124 | 0.93 |
| memory | ~ | IFOF | -3.50 | 2.20 | -1.59 | .111 | -2.13 |
| memory | ~ | ATR | 1.99 | 1.80 | 1.11 | .269 | 1.23 |
| memory | ~ | CST | -0.15 | 1.10 | -0.13 | .895 | -0.09 |
| memory | ~ | FMaj | 0.46 | 1.56 | 0.30 | .766 | 0.28 |
| memory | ~ | FMin | 0.21 | 2.28 | 0.09 | .928 | 0.13 |
| memory | ~ | CG | -1.09 | 1.04 | -1.05 | .294 | -0.72 |
| memory | ~ | CH | 1.64 | 0.99 | 1.66 | .096 | 1.00 |
| memory | ~ | ILF | 0.30 | 0.95 | 0.31 | .754 | 0.19 |
